# Activity of estrogen receptor beta expressing neurons in the medial amygdala regulates preference towards receptive females in male mice

**DOI:** 10.1101/2023.03.24.534059

**Authors:** Satoshi Takenawa, Yutaro Nagasawa, Kim Go, Yoan Chérasse, Seiya Mizuno, Kazuhiro Sano, Sonoko Ogawa

## Abstract

The processing of information regarding the sex and reproductive state of conspecific individuals is critical for successful reproduction and survival in males. Generally, male mice exhibit a preference towards sexually receptive (RF) over non-receptive females (XF) or gonadally intact males (IM). Previous studies suggested the involvement of estrogen receptor beta (ERβ) expressed in the medial amygdala (MeA) in male preference towards RF. To further delineate the role played by ERβ in the MeA in the neuronal network regulating male preference, we developed a new ERβ-iCre mouse line using the CRISPR-Cas9 system. Fiber-photometry Ca^2+^ imaging revealed that ERβ expressing neurons in the postero-dorsal part of the MeA (MeApd-ERβ^+^ neurons) were more active during social investigation towards RF compared to co-presented XF or IM mice in a preference test. Chemogenetic inhibition of MeApd-ERβ^+^ neuronal activity abolished a preference to RF in “RF vs. XF”, but not “RF vs. IM”, tests. Analysis with cre-dependent retrograde tracing viral vectors identified the principal part of the bed nucleus of stria terminalis (BNSTp) as a primary projection site of MeApd-ERβ^+^ neurons. Fiber-photometry recording in the BNSTp during a preference test revealed that chemogenetic inhibition of MeApd-ERβ^+^ neurons abolished differential neuronal activity of BNSTp cells as well as a preference to RF against XF but not against IM mice. Collectively, these findings demonstrate for the first time that MeApd-ERβ^+^ neuronal activity is required for expression of receptivity-based preference (i.e., RF vs XF) but not sex-based preference (i.e., RF vs IM) in male mice.

**Significance Statement:** In this study, by introducing a new Cre mice line for ERβ^+^ cells, we described the function of MeApd-ERβ^+^ neurons and characteristics of their neuronal activity during preference tests. Using fiber photometry and DREADD techniques we have found MeApd-ERβ^+^ neurons have a specific role in receptivity-based (receptive female vs. non-receptive female) preference but not in sex-based (receptive female vs. intact male) preference in male mice. We have also described this specific role of MeApd-ERβ^+^ neurons is achieved by regulating the neuronal activity of downstream BNSTp neurons during receptivity-based, but not sex-based, preference tests. Our findings contribute to a better understating of the function of estrogen receptor expressing neurons in the neuronal network for the male-typical reproductive behaviors.

## Introduction

The processing of information regarding the sex and reproductive state of conspecific individuals is essential for successful reproduction and survival in male rodents. Male rats and mice primarily rely on olfactory cues to discriminate and identify the right individual for the efficient expression of sexual behavior. Generally, males exhibit a preference for receptive females (RF) over non-RF (XF) or males. During a preference test with a pair of “stimulus” animals, they spend more time investigating females than males, and females in estrus more than those in non-estrus states (1–4). The action of gonadal steroid hormones is critical in preference towards sexually receptive females, as evidenced by the disruption caused by castration (1, 3, 5, 6).

Information regarding sex and reproductive status is initially processed in the main and accessory olfactory bulbs under the influence of gonadal steroids (7–11). This information is then processed in the amygdala to express preferences. Lesions of the amygdala in male rodents alter their preference for females over males (12–15). Among the different subregions of the amygdala, the medial amygdala (MeA) plays a central role in processing olfactory information that leads to the adaptive expression of reproductive behaviors (9, 16–18). Neurons in the MeA, particularly those in the posteroventral and posterodorsal (MeApd) subdivisions, project to various brain areas, such as the bed nucleus of the stria terminalis (BNST) and the ventromedial nucleus of the hypothalamus, which are involved in the expression of sexual and aggressive behaviors (19–21). Moreover, distinct neuronal subpopulations within the MeA that respond specifically to certain olfactory cues, such as males, females, and pups, have been reported (22, 23).

It is well established that testosterone regulates the expression of male-typical social behavior by acting not only on androgen receptors but also on estrogen receptors (ER), namely ERα and ERβ, after being converted to estradiol in the brain. In the MeA, both ERα and ERβ, and aromatase, which converts testosterone to estradiol, are widely expressed (24–31). Previous studies with RNAi-mediated brain site-specific knockdown in adult male mice revealed that neither type of ER in the MeA is required for the expression of sexual or aggressive behaviors (4, 32). However, the presence of ERβ in the MeA was necessary for male mice to show a preference for RFs. Importantly, the disruption of behavioral phenotype induced by ERβ silencing in the MeA was only observed when male mice were tested for preference between the RF and XF mouse, but not when they were tested for preference between an RF mouse and a gonadally intact male (IM) mouse (4). Collectively, these studies indicate that the ERβ-expressing neurons in the MeA regulate “receptivity-based preference” (i.e., RF over XF) but not “sex-based preference” (i.e., RF over IM) in male mice. However, the exact mechanism, particularly whether and how neuronal activity of ERβ-expressing neurons in the MeA is involved in this regulation, remains unknown to date.

To investigate this mechanism, we first generated a transgenic mouse line by inserting improved cyclization recombinase (iCre) just before the stop codon of the endogenous ERβ gene (*Esr2*). This allowed us to target ERβ-expressing cells using adeno-associated virus (AAV)- mediated gene delivery. Using this ERβ-iCre mouse line, we then recorded the neuronal activity of ERβ-positive neurons during preference tests in the MeApd, the primary region where ERβ-positive neurons are localized (33). Afterward, we examined whether the manipulation of MeApd-ERβ^+^ neuronal activity, using optogenetic and chemogenetic methods, would alter the preference of male mice. Furthermore, we evaluated the effect of chemogenetic inhibition of MeApd-ERβ^+^ neurons on both receptivity and sex-based preferences and the neuronal activity of the BNST, a primary downstream brain region of the MeA.

## Results

### Study 1: Fiber photometry recording of MeApd-ERβ^+^ neuronal activity during preference tests

In this study, we aimed to examine the neuronal activity of MeApd-ERβ^+^ neurons during a 10-minute preference test in freely behaving male mice. To achieve this, we used newly generated ERβ-iCre male mice that were injected with a Cre-dependent genetically encoded Ca^2+^ sensor, GCaMP7f, in the MeApd and subjected to fiber photometry recordings (Fig. 1A and 1B) during the preference tests (Fig. 1C)

**Figure 1.**
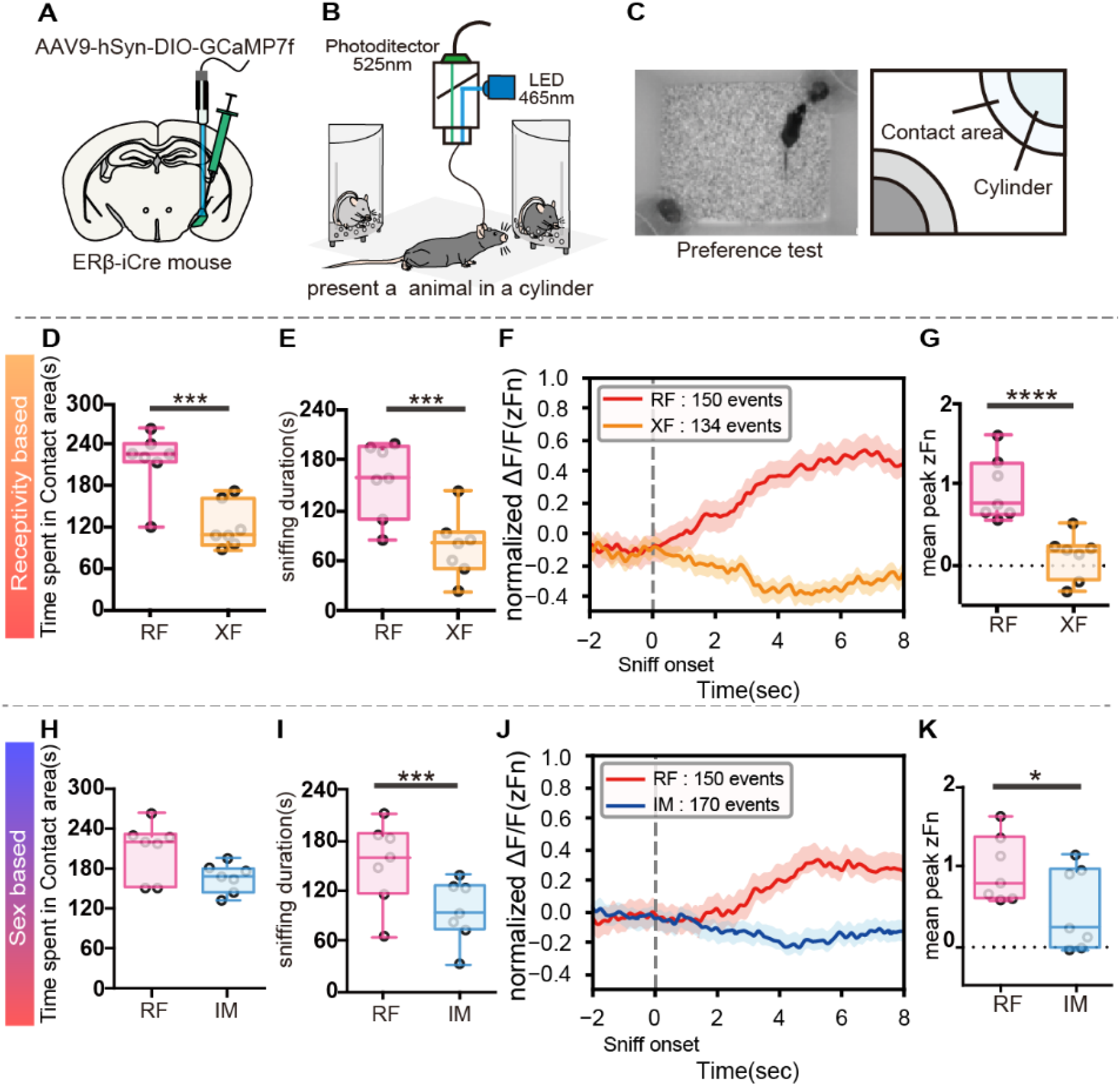
Fiber photometry recordings from MeApd-ERβ^+^ neurons during the sexual preference test. (A) A schematic diagram showing the AAV injection and fiber implantation in the MeApd. (B) A schematic diagram of the fiber-photometry system used in this study (465 nm LED emission light, filter 525 nm). (C) Representative photograph (left) and schematic diagram of preference test setup (right). (D) Duration of time (seconds) spent in contact area during RF (red) vs. XF (orange) preference tests (Mean ± SEM, n = 7, ***: *p* = 0.0004, Student’s t-test). (E) Sniffing duration (seconds) towards each stimulus during RF (red) vs. XF (orange) preference tests (Mean ± SEM, n = 7, ***: *p* = 0.0008, Student’s t-test). (F) The mean GCaMP7f signals (Z-score) during sniffing toward RF (red: 150 events) and XF (orange: 134 events) stimulus mice. The data were derived from seven male mice and shown for 10 seconds (2 seconds before to 8 seconds after) around the onset of each sniffing event. (G) The mean peak GCaMP7f signals recorded from MeApd-ERβ^+^ neurons during each sniffing event in RF (red) vs. XF (orange) preference tests. The peak value represents the maximum Z-score change within 8 seconds of sniffing onset. Activity data from 2 seconds before the onset of sniffing was used as a baseline value (Mean ± SEM, n = 7, ****: *p* <0.00001, Student’s t-test). (H) Duration (seconds) of time spent in the contact area during RF (red) vs. IM (blue) preference tests (Mean ± SEM, n = 7, *p* = 0.0550, Student’s *t* test). (I) Sniffing duration (seconds) towards each stimulus during RF (red) vs. IM (blue) preference tests (Mean ± SEM, n = 7, ***: *p* = 0.0057, Student’s t-test). (J) The mean GCaMP7f signals (Z-score) during sniffing toward RF (red: 150 events) and IM (blue: 170 events) stimulus mice. The data were derived from seven male mice and shown for 10 seconds (2 seconds before to 8 seconds after) around the onset of each sniffing event. (K) The mean peak GCaMP7f signals recorded from MeApd-ERβ^+^ neurons during each sniffing event in the RF (red) vs. IM (blue) preference tests. The peak value represents the maximum Z-score change within 8 seconds of sniffing onset. Activity data from 2 seconds before the onset of sniffing was used as a baseline value (Mean ± SEM, n = 7, *: *p* = 0.0104, Student’s t-test). MeA, medial amygdala; MeApd, MeA-posterodorsal; ER, estrogen receptor; AAV, adeno-associated virus; LED, light-emitting diode; IM, intact male; RF, receptive female; XF, non-receptive female; SEM, standard error of the mean

During receptivity-based preference tests with RF vs. XF stimuli, male mice showed a preference towards RF, as well as higher neuronal activity of MeApd-ERβ^+^ neurons during the RF investigation. Male mice spent a significantly longer time (Student’s t-test, n=7, *t* =5.428 *df*=6 *p*=0.0008***; Fig. 1D) and had a longer duration of sniffing (Student’s t-test, n=7, *t* =5.428 df=6 *p* =0.0008***; Fig. 1E), in the contact area with RF mice than with XF mice, suggesting a preference for RF mice. Additionally, fiber photometry recordings showed that MeApd-ERβ^+^ GCaMP7f signals were significantly higher during RF sniffing than XF sniffing (Fig. 1F), confirming the correlation between higher neuronal activity and preference towards RF. Further statistical analysis of neuronal activity for the entire 8 seconds (Student’s t-test, n = 7, *t* = 8.293 *df* = 6 *p*<0.0001***; Fig. 1G) confirmed that the peak value was significantly higher during sniffing towards the RF than the XF.

Similarly, during sex-based preference tests with RF vs. IM stimuli, male mice showed a preference for RF and higher MeApd-ERβ^+^ neuronal activity during RF investigation, compared to IM. Male mice tended to spend a longer time (Student’s t-test, n = 7, *t* = 1.874 *df* = 6 *p* = 0.0550; Fig. 1H) and showed a longer duration of sniffing (Student’s t-test, n = 7, *t* = 3.591 *df* = 6 *p* = 0.0057***; Fig. 1I), suggesting a preference toward RF mice. Analysis of fiber photometry recordings showed that MeApd-ERβ^+^ GCaMP7f signals during RF sniffing were significantly higher than those during IM sniffing (Fig. 1J). Statistical analysis of neuronal activity for 8 seconds confirmed that the peak value was higher during sniffing towards the RF than IM (Student’s t-test, n = 7, *t* = 3.114 *df* = 6 *p* = 0.0104*; Fig. 1K).

### Study 2: Responses of MeApd-ERβ^+^ neurons to individually presented social stimuli

In Study 1, we showed that MeApd-ERβ^+^ neurons in male mice were more active during the sniffing of RF mice compared to the sniffing of co-presented XF or IM mice. Based on these results, we could assume that these neurons specifically respond to RFs or simply respond to preferred animals. To further understand the function of MeApd-ERβ^+^ neurons, we examined how these neurons respond to stimuli (RF, XF, and IM) presented individually.

During preference tests for RF vs. Empty, male mice tended to spend more time in the RF contact area compared to the empty cylinder area (Student’s t-test n=5, *t* =2.009 df=4 *p* =0.0576; Fig. 2A). GCaMP7f signal patterns were different between the RF and empty cylinders (Fig. 2B), and peak GCaMP7f signals were much higher during the sniffing of the RF than that of an empty cylinder (Student’s t-test n=5, *t* =3.511 df=4 *p* =0.0123*; Fig. 2C).

**Figure 2.**
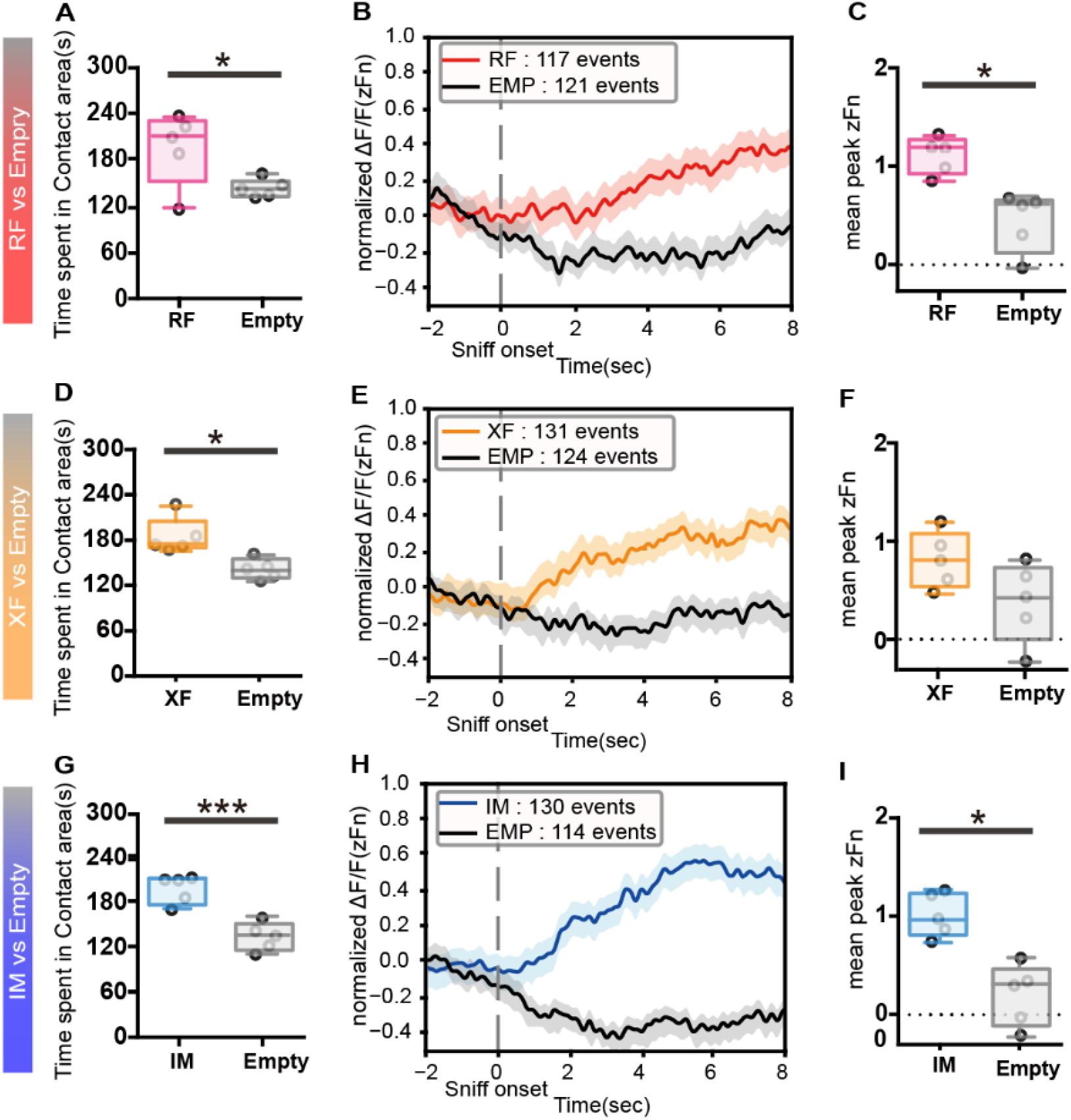
Fiber photometry recordings from MeApd-ERβ^+^ neurons in response to individually presented social stimuli. (A) Duration of time spent in the contact area during RF (red) vs. Empty (gray) preference tests (Mean ± SEM, n = 5, *p* = 0.0576, Student’s t-test). (B) The mean GCaMP7f signals (Z-score) during sniffing toward RF (red: 117 events) and Empty (gray: 121 events) cylinders. The data were derived from five male mice and shown for 10 seconds (2 seconds before to 8 seconds after) around the onset of each sniffing event. (C) The mean peak GCaMP7f signals recorded from MeApd-ERβ^+^ neurons during each sniffing event in RF (red) vs. Empty (gray) preference tests (Mean ± SEM, n = 5, *: *p* = 0.0123 Student’s t-test). (D) Duration of time (seconds) spent in the contact area during XF (orange) vs. Empty (gray) preference tests (Mean ± SEM, n = 5, *: *p* = 0.0127, Student’s t-test). (E) The mean GCaMP7f signals (Z-score) during sniffing toward XF (orange: 131 events) and Empty (gray: 124 events) cylinders. The data were derived from five male mice and shown for 10 seconds (2 seconds before to 8 seconds after) around the onset of each sniffing event. (F) The mean peak GCaMP7f signals recorded from MeApd-ERβ^+^ neurons during each sniffing event in the XF (orange) vs. Empty (gray) tests (Mean ± SEM, n = 5, *p* = 0.0951, Student’s t-test). (G) Duration of time spent in the contact area (seconds) during IM (blue) vs. Empty (gray) preference tests (Mean ± SEM, n = 5, ***: *p* = 0.0005, Student’s t-test). (H) The mean GCaMP7f signals (Z-score) during sniffing toward IM (blue: 130 events) and Empty (gray: 114 events) cylinders. The data were derived from five male mice and shown for 10 seconds (2 seconds before to 8 seconds after) around the onset of each sniffing event. (I) The mean peak GCaMP7f signals recorded from MeApd-ERβ^+^ neurons during each sniffing event in the IM (blue) vs. Empty (gray) preference tests (Mean ± SEM, *, n = 5, *p* = 0.0123, Student’s t-test). MeA, medial amygdala; MeApd, MeA-posterodorsal; ER, estrogen receptor; RF, receptive female; XF, non-receptive female; SEM, standard error of the mean

In XF vs. Empty preference tests, male mice spent more time in the contact area of the XF than the empty cylinder (Student’s t-test, n = 5; *t* = 3.480; df = 4; *p* = 0.0127*; Fig. 2D). In parallel with behavioral events, activity patterns (Fig. 2E) and peak GCaMP7f signals (Student’s t-test, n = 5; *t* = 1.576; df = 4; *p* = 0.0951; Fig. 2F) tended to be different between the sniffing of XF and empty cylinders.

Similarly, in preference tests for IM vs. Empty, male mice spent more time in the contact area of the IM (Student’s t-test, n=5, *t* =8.642 df=4 *p* =0.0005***; Fig. 2G), accompanied by higher neuronal activity during sniffing toward the IM compared to an empty cylinder (Fig. 2H and 2I, Student’s t-test n=5, *t* =3.516 df=4 *p* =0.0123* for Fig. 2P).

### Study 3: DREADD inhibit MeApd-ERβ^+^ neurons, disrupting the expression of RF vs. XF but not RF vs. IM preference

To gain a deeper understanding of the role of MeApd-ERβ^+^ neurons in the expression of preference, we conducted a study to chemogenetically inhibit these neurons and examine the effects on sniffing behavior during preference tests. For this purpose, we injected a Cre-dependent inhibitory DREADD-expressing virus or control virus bilaterally into the MeApd of ERβ-iCre mice, followed by clozapine-N-oxide (CNO) injection to chemogenetically inhibit the MeApd-ERβ+ neuronal activity (Fig. 3A and 3B). We then examined the effects on preference during the RF vs. XF and RF vs. IM tests (Fig. 3C).

**Figure 3.**
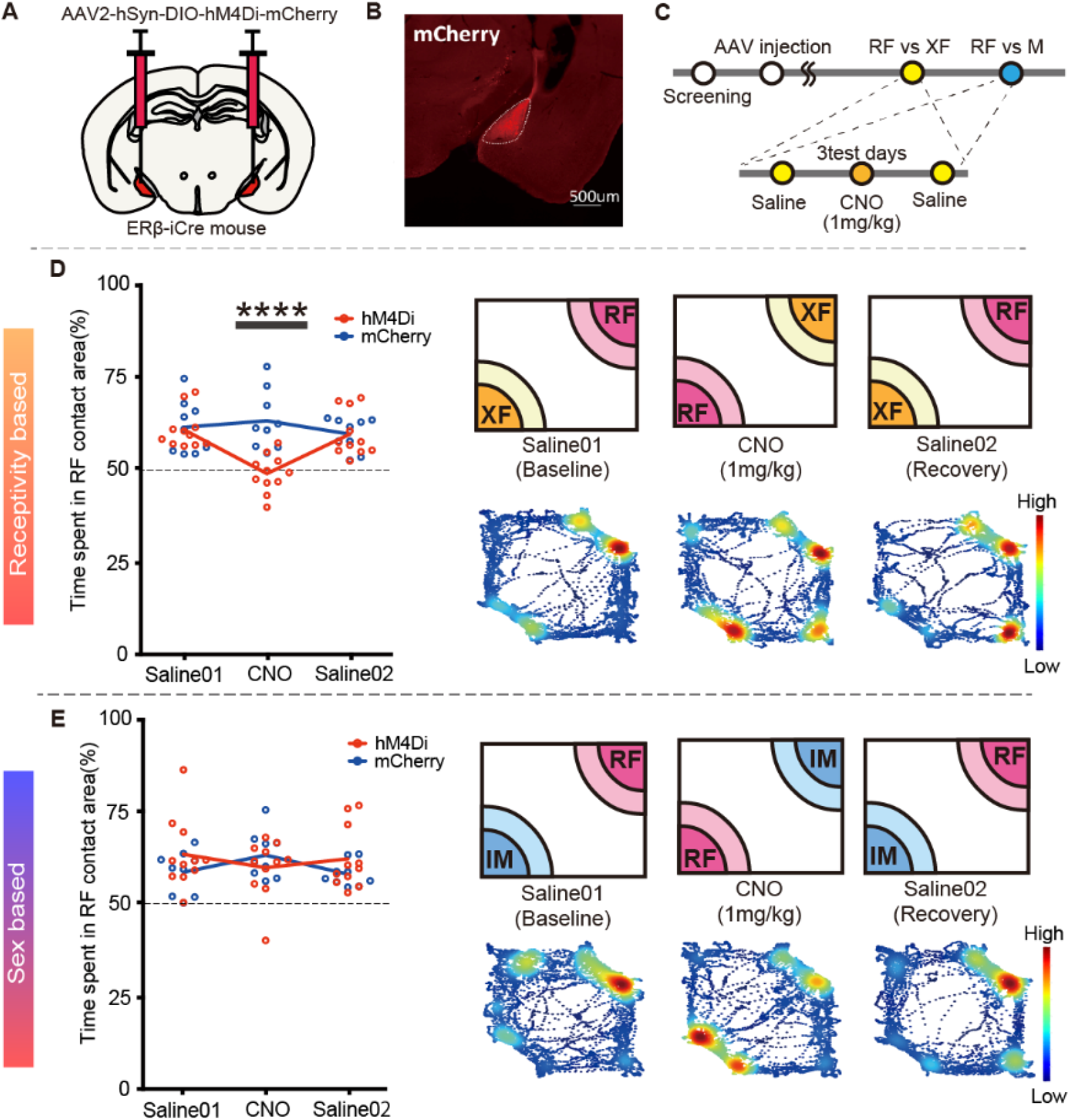
DREADD inhibition of MeApd-ERβ^+^ neurons disrupts receptivity-based, but not sex-based, preference in male mice. (A) A schematic diagram of bilateral AAV injection in the MeApd. (B) Representative image showing expression of hM4Di-mCherry in MeApd-ERβ^+^ neurons (scale = 500 μm). (C) A schematic diagram and the timeline for preference tests with DREADD inhibition. Male mice were injected with AAV and tested for receptivity and sex-based preferences on three test days (four-to five-day interval), with either saline (days 1 and 3) or CNO (day 2; 1 mg/kg) injections. (D) Right: Mean percentage (%) of time spent in the contact area of the RF cylinder during RF vs. XF tests for the hM4Di (red) and mCherry control (blue) groups. Values were obtained by dividing the RF duration by the total duration of time spent in the contact area of the RF and XF cylinders (Mean ± SEM, n = 11 for hM4Di and n = 9 for mCherry control, ****: *p* < 0.0001, Bonferroni’s test). Left: A schematic diagram of the receptivity-based (RF vs. XF) preference test and representative heat map of body position in each test performed on three consecutive days. The data were obtained from a hM4Di-injected animal. (E) Right: Mean percentage (%) of time spent in the contact area of the RF cylinder during RF vs. IM tests for the hM4Di (red) and mCherry control (blue) groups. Values were obtained by dividing the RF duration by the total duration of time spent in the contact area of RF and IM cylinders (Mean ± SEM, n = 11 for hM4Di and n = 9 for mCherry control, ns, Bonferroni’s test). Left: A schematic diagram of the sex-based (RF vs. IM) preference test and representative heat map of body position in each test performed three test days (four-to five-day interval). The data were obtained from a hM4Di-injected animal. DREADD, designer receptors exclusively activated by designer drugs; AAV, adeno-associated virus; NO, clozapine-N-oxide; MeA, medial amygdala; MeApd, MeA-posterodorsal; ER, estrogen receptor; RF, receptive female; XF, non-receptive female; SEM, standard error of the mean

In the RF vs. XF preference test, we found that CNO injection did not affect total moving distance (Fig. S2A) but clearly reduced the duration of time spent in the RF contact area (Fig. 3D) and sniffing towards the RF cylinder (Fig. S2B). Furthermore, the ratio of time spent in the RF contact area to the total duration was not significantly different from chance (repeated measures ANOVA, drug × virus *F* _(2,36)_ = 7.244 *p* = 0.0023***; Fig. 3D), indicating the abolishment of preference. In contrast to the effects seen in the RF vs. XF test, inhibition of MeApd-ERβ^+^ neuronal activity did not affect any behavioral measures in the RF vs. IM test (Fig. 3E, S2C, and S2D).

### Study 4: Role of MeApd-ERβ^+^ in the neuronal network involved in receptivity-based preference

In Study 1, MeApd-ERβ^+^ neurons showed increased activity during sniffing towards the RF cylinder in both the RF vs. XF and RF vs. IM preference tests. However, in Study 3, chemogenetic inhibition of MeApd-ERβ^+^ neuronal activity affected the expression of preference towards RF only against XF but not against IM. Based on these findings, we hypothesized that the neuronal networks activated during receptivity and sex-based preference tests may be different. To test this hypothesis, we first identified the projection patterns of MeApd-ERβ^+^ neurons by injecting a Cre-dependent anterograde viral tracer. We found that MeApd-ERβ^+^ neurons projected robustly to the BNST, particularly the principal region (BNSTp) (Fig. S3A-S3C)

To examine the effects of chemogenetic inhibition of MeApd-ERβ^+^ on the neuronal activity of BNSTp neurons, we bilaterally injected a Cre-dependent inhibitory DREADD-expressing virus to target MeApd-ERβ^+^ neurons and unilaterally injected a non-specific GCaMP7f-expressing viral vector in the BNSTp to record BNSTp neuronal activity (Fig. 4A–4C). Mice were tested four times, as shown in Fig. S4A. In RF vs. XF tests, preference for RF was disrupted in CNO-injected tests, as expected (Fig. S4B). Similar to MeApd-ERβ^+^ neurons, BNSTp neurons demonstrated higher neuronal activity toward RF during RF vs. XF preference tests with saline injection (i.e., without inhibition of MeApd-ERβ^+^ neurons; Fig. 4D). When we inhibited MeApd-ERβ^+^ neurons with CNO injection, neuronal activity of downstream BNSTp neurons was not different between RF and XF sniffing events (Fig. 4E). This was consistent with the behavioral effect of DREADD inhibition of MeApd-ERβ^+^ neurons (shown in Fig. S4B). During preference tests with CNO, changes in BNSTp GCaMP7f signals (Fig. 4E) and peak values (repeated measures ANOVA, drug × stimulus *F* _(1,20)_ = 23.80 *p* < 0.0001***; post-hoc Bonferroni’s test (RF-XF × drug); saline: p<0.0007***, CNO: *p* value>0.9999 Fig. 4F) did not differ between RF and XF sniffing events.

**Figure 4.**
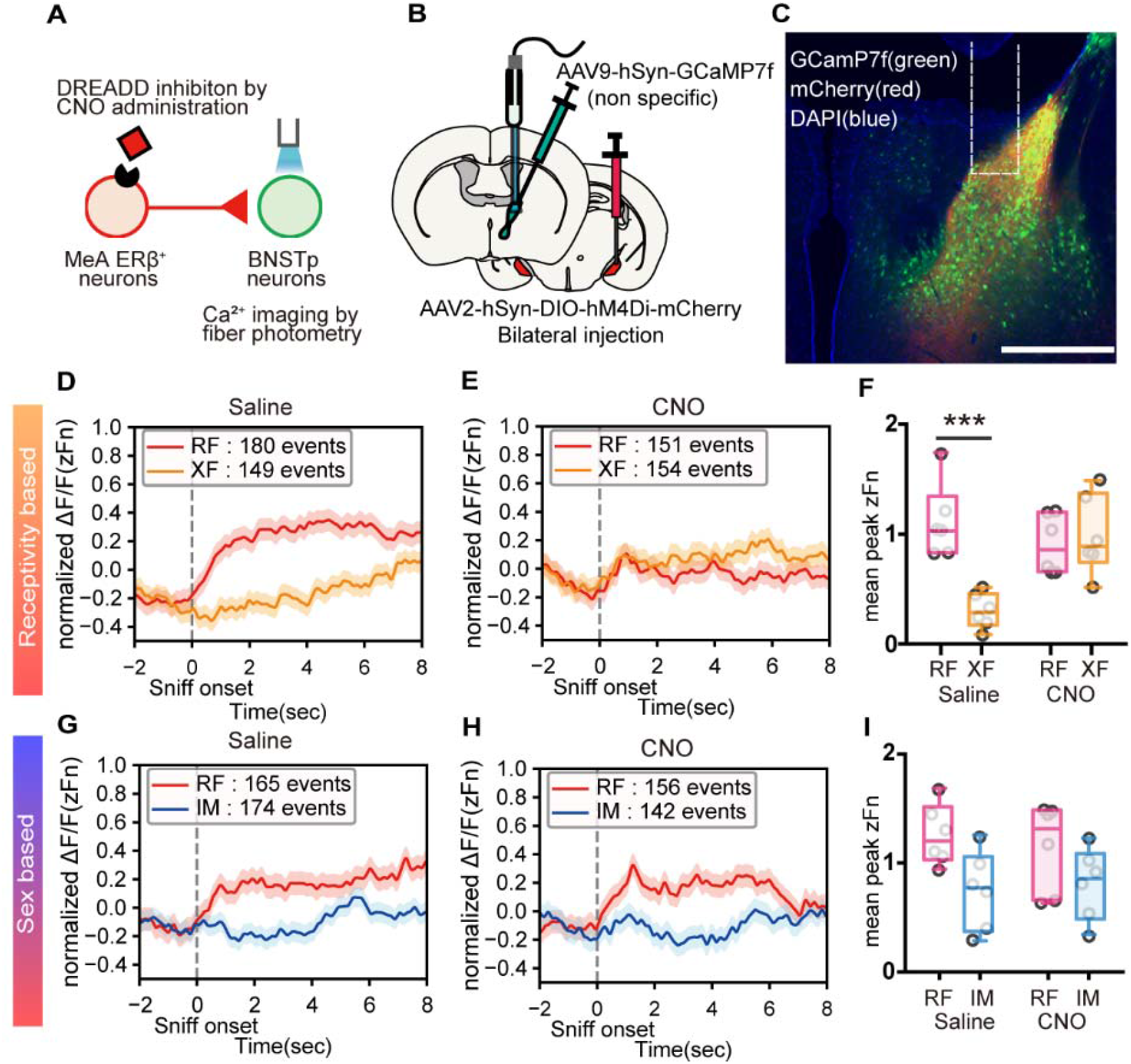
DREADD inhibition of MeApd-ERβ^+^ neurons affects BNSTp neuronal activity. (A) A schematic diagram of the experimental paradigm, in which BNSTp neuronal activity was non-specifically recorded using a fiber photometry imaging system while MeApd-ERβ^+^ neurons were inhibited. (B) A schematic diagram of the AAV injection process. A non-specific GCaMP7f-expressing virus was injected unilaterally in the BNST, and a Cre-dependent hM4Di-expressing virus was injected bilaterally in the MeApd. (C) Representative image of the BNSTp-expressing GCaMP7f (green), projecting fibers from hM4Di-expressing MeApd-ERβ^+^ neurons (red), and DAPI (blue) in the MeApd (scale bar = 500 μm). The white dots represent tracks from the inserted fiber. (D) The mean GCaMP7f signals (Z-score) during sniffing toward RF (red: 180 events) and XF (orange: 149 events) stimulus mice in saline-injected control sessions. The data were derived from six male mice and shown for 10 seconds (2 seconds before to 8 seconds after) around the onset of each sniffing event. (E) The mean GCaMP7f signals (Z-score) during sniffing toward RF (red: 151 events) and XF (orange: 154 events) stimulus mice in CNO-injected sessions. The data were derived from six male mice and shown for 10 seconds (2 seconds before to 8 seconds after) around the onset of each sniffing event. (F) The mean peak GCaMP7f signals recorded from BNSTp neurons during each sniffing event in the RF (red) and XF (orange) preference tests (Mean ± SEM, n = 6, ***: *p* < 0.001, Bonferroni’s test). (G) The mean GCaMP7f signals (Z-score) during sniffing toward RF (red: 165 events) and IM (blue: 174 events) stimulus mice in saline-injected control sessions. The data were derived from six male mice and shown for 10 seconds (2 seconds before to 8 seconds after) around the onset of each sniffing event. (H) The mean GCaMP7f signals (Z-score) during sniffing toward RF (red: 156 events) and IM (blue: 142 events) stimulus mice in CNO-injected sessions. The data were derived from six male mice and shown for 10 seconds (2 seconds before to 8 seconds after) around the onset of each sniffing event. (I) The mean peak GCaMP7f signals recorded from BNSTp neurons during each sniffing event in the RF (red) and IM (blue) preference tests (Mean ± SEM, n = 6, ***: *p* < 0.001, Bonferroni’s test). DREADD, designer receptors exclusively activated by designer drugs; AAV, adeno-associated virus; BNST, bed nucleus of the stria terminalis; BNSTp, BNST-principal; CNO, clozapine-N-oxide; DAPI, 4’,6-diamidino-2-phenylindole; MeA, medial amygdala; MeApd, MeA-posterodorsal; ER, estrogen receptor; RF, receptive female; XF, non-receptive female; SEM, standard error of the mean; IM, intact male

In contrast, inhibition of MeApd-ERβ^+^ neurons did not affect preference in RF vs. IM tests, as expected based on the findings in Study 3. The mice spent a longer time in the RF contact area than in the IM contact area under both saline and CNO injection conditions (Fig. S4E). In parallel, DREADD inhibition of MeApd-ERβ^+^ neurons did not affect BNSTp GCaMP7f signals. In both the saline and CNO conditions, changes in BNSTp GCaMP7f signals (Fig. 4G and 4H, respectively) and peak values (repeated measures ANOVA, drug × stimulus *F* _(1,20)_ = 0.3787 *p* = 0.5452 *ns*; Fig. 4I) were different between RF and IM sniffing events.

## Discussion

In this study, we investigated the possible neuronal mechanisms underlying sexual preference in male mice, with a focus on the neuronal activity of MeApd-ERβ^+^ cells. In our previous gene silencing study using RNAi methods, we demonstrated that gene expression of ERβ in the MeA is necessary for the preference of male mice toward RF over XF, but not over IM mice (4). In this study, we show that not only gene expression of ERβ but also neuronal excitation of ERβ-expressing cells, specifically in the MeApd, is critical for receptivity-based (RF vs. XF) preference but not sex-based (RF vs. IM) preference. Although previous studies in mice and rats have suggested that testosterone is necessary for males to prefer RF over XF and IM stimuli (1, 3, 6), the exact mechanism of action has not been determined. Our findings provide essential evidence for understanding the ERβ-mediated gonadal steroid action and its expressing neurons regulating male sexual preference.

### MeApd-ERβ^+^ neurons were activated in response to the sniffing of RFs

In the fiber-photometry recordings, we found the activity of MeApd ERβ^+^ neurons increased during RF investigation in both RF vs. XF and RF vs. IM preference tests, in parallel with their overall higher sniffing behavior toward RF (Study 1, Fig. 1), suggesting that the neuronal activity of MeApd-ERβ^+^ may be related to the behavioral preference of males towards RF. However, when stimulus mice were presented individually, we detected similar patterns of MeApd-ERβ^+^ activity during sniffing of all three types of stimuli, i.e., RF, XF, and IM (Study 2, Fig. 2). These findings suggest that MeApd-ERβ^+^ neurons can be similarly excited by the sniffing of RF, XF, and IM but become distinctively more active during the sniffing of RFs presented as a pair with XF or IM.

Previous studies have reported the existence of distinctive neuronal subpopulations in the MeA that respond to a specific type of stimulus, such as males, females, or pups (22), but each type of stimulus was presented individually. In contrast, our study revealed that MeApd-ERβ^+^ neurons respond differently depending on how the stimuli are presented. We conclude that the neuronal activity of MeApd-ERβ^+^ is not specific to a particular type of stimulus but underlies a preference for a specific stimulus (i.e., RF) over others (i.e., XF or IM). Our optogenetic studies further supported these findings (see Fig. S4). By optogenetically stimulating the MeApd-ERβ^+^ neurons when males started sniffing the XF mouse in RF vs. XF preference tests, we could successfully switch their preference from RF to the originally less preferred XF. These findings show artificial activation of MeApd-ERβ^+^ neurons can promote further sniffing in response to that specific stimulus (Fig. S4H).

### Inhibition of MeA ERβ^+^ neuronal activity abolished receptivity-based preference (RF vs. XF) but not sex-based preference (RF vs. IM) in male mice

Although our findings revealed that MeApd-ERβ^+^ neurons were significantly more active during the sniffing of RFs in both RF vs. XF and RF vs. IM preference tests, the effects of chemogenetic silencing of MeApd-ERβ^+^ neuronal activity on the preference behavior of male mice differed between the two tests (Study 3, Fig. 3). The preferential sniffing toward RF over XF was completely abolished by CNO injection in the hM4Di-injected group. On the other hand, males continued to exhibit more sniffing toward RF over IM even when MeApd-ERβ^+^ neurons were not activated. These findings suggest that while MeApd-ERβ^+^ neurons are active during the investigation of RF mice in both preference tests, actual activation is only necessary for receptivity-based preference but not for sex-based preference.

Our results are consistent with a previous study on site-specific ERβ silencing in the brain. The knockdown of ERβ gene expression in the MeA abolished receptivity-based preference but not sex-based preference (4). Therefore, not only the expression of ERβ but the actual neuronal activation of MeApd-ERβ^+^ neurons is necessary for receptivity-based preference in male mice.

ERβ knockdown may cause a decrease in neuronal activity among MeApd-ERβ^+^ neurons, partly through rapid nongenomic action *via* ERβ. Several reports have demonstrated that membrane-bound ERβ can modulate neuronal activity directly under the existence of 17β-estradiol, independent of their genomic action (35–37).

### BNSTp, as a primary projection site of MeApd-ERβ^+^ neurons, was activated during the sniffing of RFs in both receptivity and sex-based preference tests

The lack of effects of chemogenetic inhibition of MeApd-ERβ^+^ neuronal activity on sex-based preference led us to hypothesize that differences in mechanisms between receptivity and sex-based preferences may be at the downstream projection sites of MeApd-ERβ^+^ neurons. To test this hypothesis, we used mice generated by mating ERβ-iCre and ERβ-red fluorescent protein (RFP) mice and found that the majority of MeApd-ERβ^+^ neurons project to the BNSTp (Fig. S5A– S5C). The distinct projection pattern of MeApd-ERβ^+^ neurons led us to further investigate the role of the neuronal circuit consisting of MeApd-ERβ^+^ and BNSTp in regulating receptivity and sex-based preferences. Our fiber photometry recordings within the BNSTp revealed higher neuronal activity during RF investigation and a higher probability of being active around the RF cylinder in both RF vs. XF and RF vs. IM preference tests (Study 4, Fig. 4). This is consistent with our previous findings in fiber photometry recordings of MeApd-ERβ^+^ neurons (Study 1, Fig. 1). Moreover, previous studies have reported that neurons in the BNSTp form a network for male social behavior that responds to a specific type of stimulus presented individually (7, 8, 38, 18, 39). To this extent, our findings provide the first clear demonstration of a differential neuronal response of the BNSTp during the investigation of RF co-presented with XF or IM mice.

### Differential role of the MeApd-ERβ^+^ and BNSTp neuronal circuits in receptivity and sex-based preferences of male mice

We aimed to determine the role of functional connectivity between MeApd-ERβ^+^ neurons and BNSTp in the receptivity and sex-based preferences of male mice by inhibiting the MeApd-ERβ^+^ neurons during the BNSTp recordings (Study 4, Fig. 4). As expected, silencing of MeApd-ERβ^+^ neurons abolished preference for RF during the RF vs. XF preference test. In parallel, BNSTp neuronal activity was influenced by the silencing of MeApd-ERβ^+^ neurons, with no significant difference in the levels of neuronal activity or the probability of activation around the presented cues between RF and XF. However, in RF vs. IM preference tests, silencing of the MeApd-ERβ^+^ neurons did not affect differential BNSTp neuronal excitation during RF and IM sniffing, as well as preferential behavior toward RF over IM. These results suggest that inputs from MeApd-ERβ^+^ neurons to BNSTp are only necessary for receptivity-based and not for sex-based preference.

In summary, our study, for the first time, demonstrated that MeApd-ERβ^+^ neuronal activity is essential in the regulation of receptivity-based but not sex-based preference in male mice (Fig. 5 Summary Schematics). This functional specificity of MeApd-ERβ^+^ neurons is exerted by influencing the neuronal activity of downstream BNSTp. In conclusion, our findings contribute to a better understanding of the mechanisms of action of gonadal steroids on the neuronal network that regulates the expression of male-type reproductive behaviors. Specifically, we show that MeApd-ERβ+ neuronal activity is required for the expression of receptivity-based, but not sexbased, preference.

**Figure 5.**
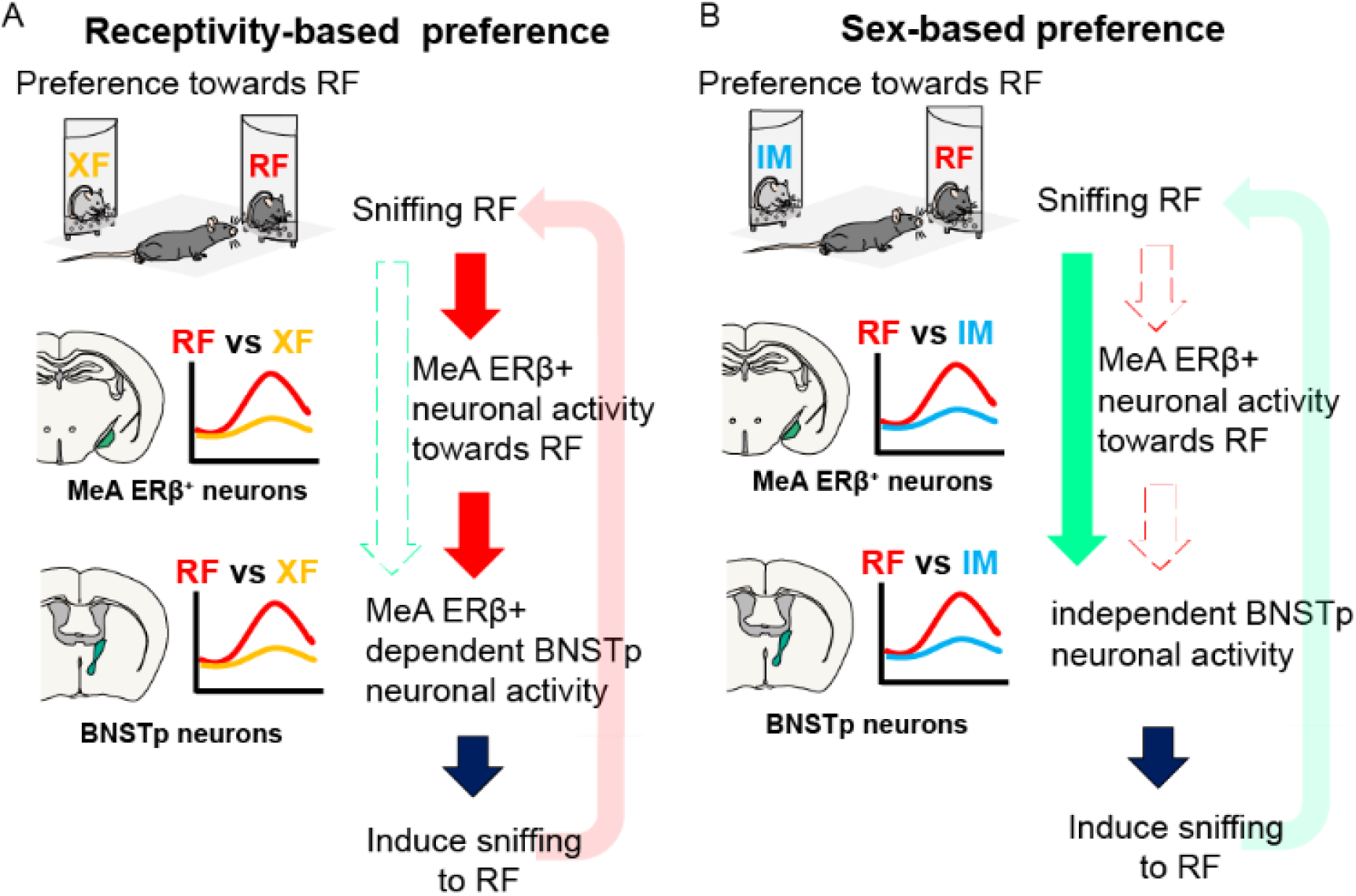
Schematic summary of MeApd-ERβ^+^ neuron function. (A) A schematic diagram illustrating the function of MeApd-ERβ^+^ neurons during receptivity-based (RF vs. XF) preference tests. Sniffing toward RF induces excitation of MeApd-ERβ^+^ neurons, which further promotes the approach and sniffing towards RF. The neuronal activity of MeApd-ERβ^+^ neurons, in turn, affects downstream BNSTp neuronal activity during a receptivity-based preference test, fully promoting receptivity-based preference. (B) A schematic diagram illustrating the function of MeApd-ERβ^+^ neurons during sex-based (RF vs. IM) preference tests. Although MeApd-ERβ^+^ neurons respond more to RF in the RF vs. IM preference test, MeApd-ERβ^+^ neuronal activity, per se, is not necessary for the expression of preference for RF over IM. Differential responses of BNSTp neurons during sniffing of RF and IM are maintained without input signals from MeApd-ERβ^+^ neurons. BNST, bed nucleus of the stria terminalis; BNSTp, BNST-principal; MeA, medial amygdala; MeApd, MeA-posterodorsal; ER, estrogen receptor; RF, receptive female; XF, non-receptive female; IM, intact male

## Materials and Methods

### Animals

We generated ERβ-iCre mice using the CRISPR-Cas9 system, as previously described (refer to Supplementary Material text, Fig. S6 and Table S1 for details) (40). Male ERβ-iCre-positive mice with a C57BL/6N background, aged between 10 and 14 weeks at the beginning of each study, were used as experimental animals. Male and female ERβ-iCre-negative mice, aged between 10 and 14 weeks, were used as stimulus animals for preference behavioral tests. All mice were housed under a 12-hour light-dark cycle, with lights turned off at noon, and were provided with food and water available *ad libitum*. All experiments were approved by the Animal Care and Use Committee and the Recombinant DNA Use Committee of the University of Tsukuba and were conducted following the National Institute of Health guidelines. All efforts were made to minimize the number of animals used and their suffering. All experimental animals were stereotaxically injected with various types of viruses under inhalation anesthesia with 1–3% isoflurane (Pfizer). The viruses used in this study are listed in Table S2.

### Preference test

The preference test apparatus and paradigms were designed based on previous studies conducted in our laboratory (4, 41). In brief, each experimental mouse was placed in a white plastic open field measuring 33 cm × 28 cm under red or dim light (10 lux) for 10 minutes. Animals used as stimuli were individually placed in a transparent acrylic quarter cylinder (7 cm base radius) with 13 small holes (Φ7 mm) on the rounded side. Time spent in the contact area, which was defined as the area 8 cm from the outer surface of the cylinders, as well as the cumulative number and duration of sniffing behavior, defined as nose touches at the perforated parts of the cylinders, were recorded. More detailed information on all experimental procedures is available in the Supplemental Methods section.

### Fiber photometry recording during preference tests

ERβ-iCre-positive mice were stereotaxically injected with either 600 nL of AAV9-hSyn-DIO-GCaMP7f at MeApd for Studies 1 and 2, or non-specific AAV9-hSyn-GCaMP7f at the BNSTp for Study 4. After a week, a NA.039, Φ230 μm glass optic fiber (RWD Life Science) was inserted 200 μm above the injection site. Fiber photometry recordings were taken during the preference tests under red light using a DORIC fiber photometry system (DORIC Lenses). A 465-nm light-emitting diode was used for the excitation of GCaMP7f, and a 525-nm emission light was filtered for recording.

All fiber photometry recordings were analyzed based on mouse sniffing behavior. Two seconds before and eight seconds after the onset of sniffing were extracted from the image data. These data were converted to dF/F0 (dF = 8 seconds from sniffing onset; F0 = mean signals from 2 seconds before sniffing onset) and normalized to the Z-score. We determined the highest signals within the 8-second window as peak signals, and the mean peak signal was derived for each stimulus. The processed fiber photometry data were analyzed and aligned with animal behavioral annotations derived from Behavioral Observation Research Interactive Software (BORIS) (42) and DeepLabCut (43) data using Python (ver. 3.8.1).

### Chemogenetic manipulation of MeApd-ERβ^+^ neurons

ERβ-iCre-positive mice were stereotaxically injected with either 300 μl of AAV2-hSyn-DIO-hM4Di-mCherry or AAV2-hSyn-DIO-mCherry control virus at the MeApd. This was followed by two separate studies, one involving simple behavioral analysis (Study 3), and the other involving combined analysis with fiber photometry recording in the BNSTp (Study 4). On the test day, animals were intraperitoneally injected with either saline or CNO (Sigma-Aldrich) at a dose of 1 mg/kg body weight, 15 minutes before the test.

## Statistical analysis

Statistical analyses were performed using GraphPad Prism 9, with all data analyzed by Student’s t-test or analysis of variance. Differences were considered significant at *p* <0.05*, *p* <0.01**, *p* <0.001***, and *p* <0.0001****. The details are summarized in Table S3.

## Acknowledgments

The authors thank Dr. C. Pavlides for critical reading of the manuscript. We also thank prof. Aki Takahashi for technical advice. Y. Asano and A. Sagehashi for their technical and secretarial assistance. This work was supported by grant-in-aid for Scientific Research 15H05724 and 22H02941 to SO.

## References

1. E. Rose, L. C. Drickamer, Castration, sexual experience, and female urine odor preferences in adult BDF1 male mice. Bull. Psychon. Soc. 5, 84–86 (1975).

2. M. de Bruijn, M. Broekman, P. van der Schoot, Sexual interactions between estrous female rats and castrated male rats treated with testosterone propionate or estradiol benzoate. Physiol. Behav. 43, 35–39 (1988).

3. K. Xiao, Y. Kondo, Y. Sakuma, Sex-specific effects of gonadal steroids on conspecific odor preference in the rat. Horm. Behav. 46, 356–361 (2004).

4. M. Nakata, et al., Effects of prepubertal or adult site-specific knockdown of estrogen receptor β in the medial preoptic area and medial amygdala on social behaviors in male mice. eNeuro 3, ENEURO.0155-15.2016 (2016).

5. S. Dhungel, S. Urakawa, Y. Kondo, Y. Sakuma, Olfactory preference in the male rat depends on multiple chemosensory inputs converging on the preoptic area. Horm. Behav. 59, 193–199 (2011).

6. Y. Kondo, H. Hayashi, Neural and hormonal basis of opposite-sex preference by chemosensory signals. Int. J. Mol. Sci. 22, 8311 (2021).

7. J. Bakker, M. J. Baum, A. K. Slob, Neonatal inhibition of brain estrogen synthesis alters adult neural Fos responses to mating and pheromonal stimulation in the male rat. Neuroscience 74, 251–260 (1996).

8. J. M. Fiber, J. M. Swann, Testosterone differentially influences sex-specific pheromone-stimulated Fos expression in limbic regions of Syrian hamsters. Horm. Behav. 30, 455–473 (1996).

9. K. L. Martel, M. J. Baum, A centrifugal pathway to the mouse accessory olfactory bulb from the medial amygdala conveys gender-specific volatile pheromonal signals. Eur. J. Neurosci. 29, 368–376 (2009).

10. M. Yoshikage, I. Toshiaki, K. Seiichi, K. Nobuo, M. Nishimura, Sex steroids modulate the signals from volatile female odors in the accessory olfactory bulb of male mice. Neurosci. Lett. 413, 11–15 (2007).

11. T. Inbar, R. Davis, J. F. Bergan, A sex-specific feedback projection from aromatase-expressing neurons in the medial amygdala to the accessory olfactory bulb. J. Comp. Neurol. 530, 648–655 (2022).

12. Y. Kondo, Lesions of the medial amygdala produce severe impairment of copulatory behavior in sexually inexperienced male rats. Physiol. Behav. 51, 939–943 (1992).

13. Y. Kondo, B. D. Sachs, Disparate effects of small medial amygdala lesions on noncontact erection, copulation, and partner preference. Physiol. Behav. 76, 443–447 (2002).

14. P. M. Maras, A. Petrulis, Chemosensory and steroid-responsive regions of the medial amygdala regulate distinct aspects of opposite-sex odor preference in male Syrian hamsters. Eur. J. Neurosci. 24, 3541–3552 (2006).

15. A. Petrulis, Neural mechanisms of individual and sexual recognition in Syrian hamsters (Mesocricetus auratus). Behav. Brain Res. 200, 260–267 (2009).

16. M. J. Baum, Sexual differentiation of pheromone processing: Links to male-typical mating behavior and partner preference. Horm. Behav. 55, 579–588 (2009).

17. K. Hashikawa, Y. Hashikawa, A. Falkner, D. Lin, The neural circuits of mating and fighting in male mice. Curr. Opin. Neurobiol. 38, 27–37 (2016).

18. D. W. Bayless, N. M. Shah, Genetic dissection of neural circuits underlying sexually dimorphic social behaviours. Philos. Trans. R. Soc. Lond. B. Biol. Sci. 371, 20150109 (2016).

19. N. S. Canteras, R. B. Simerly, L. W. Swanson, Organization of projections from the medial nucleus of the amygdala: A PHAL study in the rat. J. Comp. Neurol. 360, 213–245 (1995).

20. H.-W. Dong, G. D. Petrovich, L. W. Swanson, Topography of projections from amygdala to bed nuclei of the stria terminalis. Brain Res. Rev. 38, 192–246 (2001).

21. D. J. Anderson, Optogenetics, Sex, and Violence in the Brain: Implications for Psychiatry. Biol. Psychiatry 71, 1081–1089 (2012).

22. Y. Li, et al., Neuronal representation of social information in the medial amygdala of awake behaving mice. Cell 171, 1176–1190.e17 (2017).

23. S. Yao, J. Bergan, A. Lanjuin, C. Dulac, Oxytocin signaling in the medial amygdala is required for sex discrimination of social cues. eLife 6, e31373 (2017).

24. A. Foidart, N. Harada, J. Balthazart, Aromatase-immunoreactive cells are present in mouse brain areas that are known to express high levels of aromatase activity. Cell Tissue Res. 280, 561–574 (1995).

25. P. J. Shughrue, M. V. Lane, I. Merchenthaler, Comparative distribution of estrogen receptor-α and -β mRNA in the rat central nervous system. J. Comp. Neurol. 388, 507–525 (1997).

26. P. Shughrue, P. Scrimo, M. Lane, R. Askew, I. Merchenthaler, The distribution of estrogen receptor-β mRNA in forebrain regions of the estrogen receptor-α knockout mouse. Endocrinology 138, 5649–5652 (1997).

27. S. Ogawa, et al., Survival of reproductive behaviors in estrogen receptor β gene-deficient (βERKO) male and female mice. Proc. Natl. Acad. Sci. 96, 12887–12892 (1999).

28. S. W. Mitra, et al., Immunolocalization of Estrogen Receptor β in the Mouse Brain: Comparison with Estrogen Receptor α. Endocrinology 144, 2055–2067 (2003).

29. S. Ogawa, et al., Abolition of male sexual behaviors in mice lacking estrogen receptors α and ß (αßERKO). Proc. Natl. Acad. Sci. 97, 14737–14741 (2000).

30. R. J. Handa, S. Ogawa, J. M. Wang, A. E. Herbison, Roles for oestrogen receptor β in adult brain function. J. Neuroendocrinol. 24, 160–173 (2012).

31. K. Sano, et al., Pubertal activation of estrogen receptor α in the medial amygdala is essential for the full expression of male social behavior in mice. Proc. Natl. Acad. Sci. 113, 7632–7637 (2016).

32. K. Sano, M. C. Tsuda, S. Musatov, T. Sakamoto, S. Ogawa, Differential effects of sitespecific knockdown of estrogen receptor α in the medial amygdala, medial pre-optic area, and ventromedial nucleus of the hypothalamus on sexual and aggressive behavior of male mice. Eur. J. Neurosci. 37, 1308–1319 (2013).

33. S. Sagoshi, et al., Detection and characterization of estrogen receptor beta expression in the brain with newly developed transgenic mice. Neuroscience 438, 182–197 (2020).

34. B. L. Roth, DREADDs for Neuroscientists. Neuron 89, 683–694 (2016).

35. M. Wong, R. L. Moss, Long-term and short-term electrophysiological effects of estrogen on the synaptic properties of hippocampal CA1 neurons. J. Neurosci. 12, 3217–3225 (1992).

36. T. Smejkalova, C. S. Woolley, Estradiol acutely potentiates hippocampal excitatory synaptic transmission through a presynaptic mechanism. J. Neurosci. 30, 16137–16148 (2010).

37. J. Cao, J. Meitzen, Perinatal activation of ERα and ERβ but not GPER-1 masculinizes female rat caudate-putamen medium spiny neuron electrophysiological properties. J. Neurophysiol. 125, 2322–2338 (2021).

38. H. A. Halem, J. A. Cherry, M. J. Baum, Vomeronasal neuroepithelium and forebrain Fos responses to male pheromones in male and female mice. J. Neurobiol. 39, 249–263 (1999).

39. B. Yang, T. Karigo, D. J. Anderson, Transformations of neural representations in a social behaviour network. Nature 608, 741–749 (2022).

40. Y. Hasegawa, et al., Generation of CRISPR/Cas9-mediated bicistronic knock-in *ins1-cre* driver mice. Exp. Anim. 65, 319–327 (2016).

41. M. C. Tsuda, S. Ogawa, Long-lasting consequences of neonatal maternal separation on social behaviors in ovariectomized female mice. PLOS ONE 7, e33028 (2012).

42. O. Friard, M. Gamba, BORIS: a free, versatile open-source event-logging software for video/audio coding and live observations. Methods Ecol. Evol. 7, 1325–1330 (2016).

43. A. Mathis, et al., DeepLabCut: markerless pose estimation of user-defined body parts with deep learning. Nat. Neurosci. 21, 1281–1289 (2018).

