## Supplemental information for "Activity of estrogen receptor beta expressing neurons in the medial amygdala regulates preference towards receptive females in male mice"

Sonoko Ogawa, Ph.D.

ORCID : 0000-0003-3976-9987

This PDF file includes:

Supporting text

Figures S1 to S10

Tables S1 to S4

SI References

Supporting Information Text

**Supplemental Results**

**Studies 1 and 2: Plotting the firing probability of MeApd-ERβ^+^ neurons using Kernel density estimation during the preference test**

During the receptivity-based preference tests (RF vs. XF), male mice demonstrated a preference for RF stimuli, as shown in Fig. 1. Furthermore, we analyzed the fiber photometry recordings to establish a relationship between the neuronal activity (Z ≥1) of MeApd-ERβ^+^ neurons and the position of the animal (regardless of sniffing behavior). To achieve this, we used kernel density estimation (KDE) methods with Scott’s rule (1). Using KDE, we not only analyzed neuronal activity but also demonstrated the probability of these activities occurring by showing them on a two-dimensional virtual map derived from the KDE analysis. Since MeApd-ERβ^+^ neurons were not always active during the sniffing of stimulus mice, we needed to show the probability of MeApd-ERβ^+^ neurons being activated.

KDE analysis visually demonstrated MeApd-ERβ^+^ neurons had a higher probability to show enhanced neuronal activity around the RF contact area than that of XF (Fig. S1A; Student’s t-test, n = 7, *t* =5,462 *df* = 6 *p* = 0.0008*** for Fig. S1B). During sex-based preference tests with RF vs. IM stimuli, male mice showed a preference towards RF, similar to the results of RF vs. XF tests. The KDE analysis also showed that MeApd-ERβ^+^ neurons had a higher probability of exhibiting enhanced neuronal activity towards RF than IM (Fig.S1C; Student’s t-test, n = 7, t = 7.545 df = 6 *p* < 0.0001*** for Fig. S1D). These results suggest that higher neuronal activity of MeApd-ERβ^+^ cells is associated with the preference of animals for RF stimuli compared to XF or IM stimuli KDE analysis revealed a relationship between the location of the animal and neuronal activity. We successfully demonstrated graphically that MeApd-ERβ^+^ neurons are consistently active around RF in both RF vs. XF and RF vs. IM tests.

When we conducted preference tests for RF vs. Empty cylinders, KDE analysis showed that the probability of MeApd-ERβ^+^ neurons exhibiting enhanced neuronal activity was higher towards RF compared to an empty cylinder (Fig. S1E; Student’s t-test, n = 5, t = 4.693 df = 4 *p* = 0.0047* for Fig. S1F). During the preference tests for XF vs. Empty, KDE analysis showed that the probability of MeApd-ERβ^+^ neurons exhibiting enhanced neuronal activity was visually higher in the XF contact area compared to an empty cylinder, although it was not statistically significant (Fig. S1G; Student’s t-test n=5, t=1.442 df=4 ns for Fig. S1H). Similarly, in preference tests for IM vs. Empty, KDE analysis demonstrated that the probability of MeApd-ERβ^+^ neurons exhibiting enhanced neuronal activity was higher in the IM contact area compared to the empty cylinder (Fig. S1I; Student’s t-test, n = 5, t = 4.897 df = 4 *p* = 0.0040**; Fig. S1J).

**Study 3: DREADD inhibition of MeApd-ERβ^+^ neurons during preference behavior did not affect total animal movement**

During the RF vs. XF tests, we observed that inhibiting MeApd-ERβ^+^ neuronal activity through CNO injection did not have an effect on the total moving distance of the animals (Repeated measures ANOVA, Interaction (drug x virus) *F* _(2,36)_ = 0.7302 *p* = 0.4999 *ns*; Fig. S2A). However, it significantly reduced the time spent sniffing towards the RF cylinder (repeated measures ANOVA, Interaction (drug × virus) *F* _(2,36)_ = 7.196 *p* = 0.0024***; Fig. S2B). In contrast, there were no significant changes in any measures during the RF vs. IM tests even when MeApd-ERβ^+^ neuronal activity was inhibited (Fig. S2C and S2D).

**Study 4:** **Viral tracing of MeApd-ERβ^+^ neurons**

To investigate the afferent projections of MeApd-ERβ^+^ neurons, we used a Cre-dependent viral tracer (Fig. S3A), which specifically labeled the synaptic terminals of these neurons. Across sections ranging from Bregma +0.14 mm to - 3.08 mm, we found that the majority of MeApd-ERβ^+^ neurons projected to the BNST, while very few projected to the hypothalamus (Fig. S3B). Additionally, we observed that the synaptic terminals of MeApd-ERβ+ neurons were highly concentrated in the BNSTp (Fig. S3C).

**Effects of DREADD inhibition of MeApd-ERβ^+^ neurons on preference behavior and BNSTp firing probability**

To investigate the effects of chemogenetic inhibition of MeApd-ERβ^+^ neurons on the activity of BNSTp neurons, we injected a viral vector that induces GCaMP7f expression non-specifically (see Fig. 4A–4C). The mice were tested four times (Fig. S4A). In the RF vs. XF test, as expected, CNO injection disrupted preference towards RF by affecting the duration that male mice spent in the RF contact area (repeated measures ANOVA, drug vs. cue (n = 6) Interaction (drug x cue) *F* _(1,20)_ = 13.64 *p* = 0.0014**; Fig. S4B). In parallel with the effect of chemogenetic inhibition of MeApd-ERβ^+^ neurons (see Fig. 4D–4F), the KDE analysis (Fig. S4C) also revealed that the probability of BNSTp neurons exhibiting prominent neuronal activity against RF seen in the saline control condition was abolished during preference tests under CNO injection (repeated measures ANOVA, drug vs cue(n=6) Interaction (drug x cue) F (1,20) =23.80 *p* <0.0001; Fig. S4D).

In contrast, inhibition of MeApd-ERβ^+^ neurons did not affect preference in RF vs. IM tests (Fig. S4E), consistent with the findings of Study 3. Male mice spent more time in the RF contact area under both saline and CNO injection conditions (repeated measures ANOVA, drug vs. cue (n = 6) Cue *F* _(1,20)_ = 10.90 *p =* 0.0036** *ns*; Fig. S4E). The KDE analysis (Fig. S4F) also showed that the probability of the BNSTp neurons exhibiting prominent neuronal activity was higher around the RF contact area than for the IM in both saline and CNO injection conditions (repeated measures ANOVA, drug vs. cue (n = 6) Interaction (drug × cue) *F* _(1,20)_ = 0.6713 *p =* 0.4223 *ns*; Fig. S4G).

**Results for Supplemental Study: Effects of optogenetic stimulation of MeApd-ERβ^+^ neurons on the levels of sniffing behavior**

Based on the fiber photometry recordings from MeApd-ERβ^+^ neurons in Studies 1 and 2, we have hypothesized that activation of MeApd-ERβ^+^ neurons by sniffing of a specific stimulus, such as RF, may further enhance sniffing towards the same stimulus (i.e., RF), resulting in a preference for RF over XF. To test this hypothesis, we examined whether optogenetical stimulation of MeApd-ERβ^+^ neurons during sniffing towards a non-preferred stimulus (e.g., XF) might enhance sniffing to that specific stimulus.

We unilaterally injected mice with AAV9-hSyn-DIO-ChR2-EYFP (Fig. S5A) and performed a 15-minute RF vs. XF preference test while optogenetically stimulating ChR2-expressing MeApd-ERβ^+^ neurons only when the mice sniffed the XF cylinder during the middle 5-minute block (Fig. S5B). As expected, during the first (pre-stimulation) and third (post-stimulation) 5-minute blocks, the mice showed higher levels of sniffing toward RF than XF (Fig. S5C and S5D). However, during the second 5-minute block, by optogenetically stimulating MeApd-ERβ^+^ neurons, the preference was reversed (repeated measures ANOVA, Interaction (time x cue) *F* _(14,112)_ = 5.976 *p* < 0.0001*** for Fig. S5D). Further analysis of behavioral changes between the three 5-minute blocks revealed that the sniffing duration (repeated measures ANOVA, Interaction (time × cue) *F* _(2,16)_ = 18.30 *p* < 0.0001***; Fig. S5E) and the number of sniffings of the XF cylinder (repeated measures ANOVA, Interaction (time × cue) *F* _(2,16)_ = 20.89 *p* < 0.0001***; Fig. S5F) significantly increased without affecting sniffing duration per event (repeated measures ANOVA, Interaction (time × cue) *F* _(2,16)_ = 1.966 *p =* 0.1723 ns; Fig. S5G). These findings suggest that optogenetic stimulation of MeApd-ERβ^+^ neurons during the investigation of non-preferred XF could drive the animal to repeatedly investigate XF and form an artificial preference for it over RF (Fig. S5H).

**Supplemental methods**

**Generation and verification of ERβ-iCre mice**

ERβ-iCre mice were generated using the CRISPR-Cas9 system, as described previously (2). A *p2A-iCre rGpA* sequence was inserted on the 5’ side of the stop codon of the C57BL/6N mouse *Esr2* gene (Fig. S6A). All animals resulting from the CRISPR-Cas9 method were screened for successful knock-in of *Esr2*-*iCre*, followed by unintentional random integration of Cas9 expression and donor DNA vectors. The resulting lines were sequenced upstream of the *iCre* gene to confirm the absence of unintentional mutations in the *Esr2* gene. All the primers used are listed in Table S1.

Since ERβ is expressed in neuronal embryonic cells in mice (3, 4), we could not confirm the expression pattern of *iCre* by reporter mouse assays. Instead, we examined *iCre* expression in ERβ-expressing cells by backcrossing ERβ-iCre mice with previously reported ERβ-red fluorescent protein (RFP)-tg mice (5). We injected ERβ-iCre and ERβ-RFP double-positive male mice (n = 3; eight weeks old) with 300 nL of AAV2-hSyn-DIO-green fluorescent protein (GFP) in the MeApd (Fig. S6B) and perfused them two weeks later. Under a fluorescent microscope, we observed the expression of Cre-induced GFP and ERβ-RFP in MeApd (between Bregma -1.56 and -2.04) (Fig. S6C). Out of 2505 GFP-positive cells, 2229 cells (89%) were RFP-positive (Fig. S6D), indicating that Cre was successfully expressed in ERβ-positive cells in the MeApd. Detailed methods are described below.

**Animal housing and surgical procedure**

Male ERβ-iCre-positive mice with a C57BL/6N background, aged between 10 and 14 weeks at the beginning of each study, were used as experimental animals. Male and female ERβ-iCre-negative mice, aged between 10 and 14 weeks, were used as stimulus animals for preference behavioral tests. All mice were housed under a 12-hour light-dark cycle, with lights turned off at noon, and were provided with food and water available *ad libitum*. All experiments were approved by the Animal Care and Use Committee and the Recombinant DNA Use Committee of the University of Tsukuba and were conducted following the National Institute of Health guidelines. All efforts were made to minimize the number of animals used and their suffering.

All experimental animals were stereotaxically injected with various types of viruses under inhalation anesthesia with 1–3% isoflurane (Pfizer). Virus injections were performed using a Hamilton syringe 7000RN with a custom-made 33-gauge, 45° beveled needle tip connected to a Micro4-injection pump (World Precision Instruments). The injection site coordinates were determined based on the Mouse Brain in Stereotaxic Coordinates (6). Detailed information is provided for each experiment, and all viruses used in this study are listed in Table S2. After surgery, all mice were individually housed in plastic cages (12.5 x 20 x 11 cm). Two weeks after surgery, all mice were screened for baseline preference towards RF against XF using the test paradigm described below. Mice that failed to show a preference towards the RF were excluded from the study.

**A detailed description of the preference test**

The preference test apparatus and paradigms were designed based on previous studies conducted in our laboratory (7, 8). In brief, each experimental mouse was placed in a white plastic open field (33 cm × 28 cm) under red or dim light (10 lux), depending on the paradigm, with clean bedding, and tested against a pair of stimulus mice, unless otherwise described. The test duration was either 10 or 15 minutes, as stated in each study. All behavioral tests were performed starting at Zeitgeber time 15.

The stimulus mice were individually placed in a transparent acrylic quarter cylinder (7 cm in base radius, 17 cm in height) with 13 small holes (Φ7 mm) near the bottom 3 cm on the rounded side. Before being used as stimulus mice, they were habituated to the cylinders more than three times. All stimulus female mice (RF and XF) were ovariectomized for more than two weeks before testing, under inhalation anesthesia with 1-3% isoflurane, and group-housed (four mice per cage). RF mice were injected subcutaneously with 1 μg 17β-estradiol (Sigma-Aldrich) in 0.1 mL sesame oil at 48 and 24 hours and 500 μg progesterone (Sigma-Aldrich) in 0.1 mL sesame oil at 4 hours before testing. The IM stimulus mice were single-housed for more than one week before testing.

On the day of the preference tests, pairs of RF and XF mice were used in the receptivity-based test, and pairs of RF and IM mice were used in the sex-based preference test. In both tests, stimuli were presented to singly housed C57BL/6N intact tester males before being used as stimuli for the preference tests. Only pairs in which tester males preferred RF mice over XF or IM mice were used for the preference tests on that day.

At the start of the preference tests, a pair of cylinders, each containing different types of stimulus mice (RF, XR, or IM), was placed at the two diagonal corners. The placement of the cylinders was counterbalanced. For some tests, empty cylinders were also used. All preference tests were recorded with a CCD camera placed 70–120 cm above the open field, depending on the experimental setup. The test mice sniffing behavior was quantified using BORIS (Friard & Gamba, 2016) and DeepLabCut (ver. 2.1.8.2) (Mathis et al., 2018) for Studies 1, 2, and 4, and the automated Ogawa-Type Social Interaction Test System (O'HARA & Co., Ltd.) (8) for Study 3. In both cases, the time spent in the contact area, defined as the area 8 cm from the outer surface of the cylinders, and the cumulative number and duration of sniffing behaviors, defined as nose touches at the perforated parts of the cylinders, were recorded.

**Fiber photometry recording and data analysis**

ERβ-iCre positive mice were stereotaxically injected with either 600 nL of AAV9-hSyn-DIO-GCaMP7f at MeApd (unilateral, AP: -1.94 mm, ML: ±2.40 mm, DV: -4.75 mm) for Studies 1 and 2, and non-specific AAV9-hSyn-GCaMP7f at the BNSTp (unilateral, AP: -0.22 mm, ML: ±0.5 mm, DV: -3.50 mm) for Study 4. After a week of virus injection, a NA.039, Φ230 μm glass optic fiber (RWD Life Science) was inserted 200 μm above the injection site and fixed with resin dental cement (Tokuyama Dental) mixed with carbon black (Sigma-Aldrich). The mice were habituated to the test condition more than three times, including the optic cable connection, for 10 minutes at least three weeks after the last surgery. Fiber photometry recordings were performed during the preference tests under red light using a DORIC fiber photometry system (DORIC Lenses). The excitation of GCaMP7f was done using a 465-nm light-emitting diode, and a 525-nm emission light was filtered for recording. After the last recording session, all mice that were subjected to fiber photometry recordings were perfused to confirm viral infection and fiber placement, as described below.

The movements of the mouse were recorded using a DORIC camera system (DORIC Lenses) linked to a fiber photometry system. The recorded image data underwent a 4 Hz low-pass filter. Sniffing behavior was annotated using BORIS (9) while body movement was tracked using DeepLabCut (ver. 2.1.8.2) (10) in Python (ver. 3.7.6). All fiber photometry recordings were analyzed based on mouse sniffing behavior. The image data from 2 seconds before and 8 seconds after the onset of sniffing were extracted, converted to dF/F0 (dF = 8 seconds from sniffing onset, F0 = mean signals from 2 seconds before sniffing onset), and normalized to the Z-score. The peak signals were determined within the 8-second window, and the mean peak signal was determined for each stimulus. Furthermore, the GCaMP signals with Z ≥1 activity during sniffing behavior were sorted as prominent neuronal activity and further processed for analysis. The processed fiber photometry data were analyzed and aligned with animal behavioral annotations derived from BORIS and DeepLabCut data using Python (ver. 3.8.1).

For the KDE analysis shown in Fig. S1 and S4, all imaging data were filtered using a 4 Hz low-pass filter and smoothed using a 60-second moving average. The data were then converted to Z-scores, and the XY coordinates of mouse body movements were virtually plotted into a 330 x 280 grid and aligned. To generate a KDE plot based on the aligned XY data, we used the scipy.stats.gaussian_KDE package in Scipy (ver. 1.4.1) and determined the KDE bandwidth selection using Scott’s rule (1). To compare the KDE probability between neuronal activity and overall body position, we stacked data across all animals, extracted the body positions accompanied by Z ≤1 neuronal activity, and plotted the KDE on the virtual grid (KDE plot, Z ≤1). To obtain comparable data for each stimulus, we calculated the KDE score within the contact area of each stimulus and termed it the KDE score.

**Chemogenetic manipulation of MeApd-ERβ^+^ neurons**

ERβ-iCre positive mice were stereotaxically injected with either 300 μl of AAV2-hSyn-DIO-hM4Di-mCherry or an AAV2-hSyn-DIO-mCherry control virus bilaterally at the MeApd (AP: -1.94 mm; ML: ±2.40 mm; DV: -4.75 mm) to investigate the chemogenetic inhibition of MeApd- ERβ^+^ neuronal activity on preference behavior in male mice in either simple behavioral analysis (Study 3) or combined analysis with fiber photometry recording in the BNSTp (Study 4). Starting three weeks after surgery, the mice were habituated to intraperitoneal injections of 0.1 mL saline for three days. They were then treated with either saline (day 1 or 3) or CNO (Sigma-Aldrich) at a dose of 1 mg/kg BW (day 2), 15 minutes prior to the test. After the last behavioral test, the mice were perfused, and viral expression was confirmed by immunohistochemical detection of mCherry protein.

**Optogenetic stimulation of MeApd-ERβ^+^ neurons**

ERβ-iCre positive mice were stereotaxically injected with 300 μl of AAV2-EF1α-DIO-ChR2-EFYP in the right MeApd (AP: -1.94 mm; ML: ±2.40 mm; DV: -4.75 mm). After one week, an NA0.50, Φ250-μm plastic fiber was inserted 200 μm above the injection site and fixed with resin dental cement (Tokuyama Dental) mixed with carbon black (Sigma-Aldrich) on the skull. Two weeks later, the mice were habituated for 15 minutes to the test condition, including the optic cable connection, three times prior to the preference test. Optogenetic stimulation was delivered manually using a 473 nm laser (LUCIR Inc.) connected to a pulse generator in 20 Hz bursts (10 ms each). The detailed experimental design is described in Supplemental Study below. Animal sniffing behavior was annotated using BORIS (ver. 8.5) (9), and body movement was tracked using DeepLabCut (Mathis et al., 2018) (ver. 2.1.8.2) in Python (ver. 3.7.6) as described previously. Video recordings during the test sessions were analyzed and aligned with the BORIS and DeepLabCut data using Python (ver. 3.8.1). At the end of all behavioral tests, mice were perfused 90 minutes after a 5-minute optogenetic stimulation (20 seconds of 20 Hz bursts, 40 seconds interval) in their home cage, and the fiber placement was confirmed.

**Tissue preparation and immunohistochemistry**

The mice were anesthetized with pentobarbital (1 mg/kg) and heparin (1000 units/kg) and transcardially perfused with phosphate-buffered saline followed by 0.1 M phosphate buffer (PB) containing 4% paraformaldehyde. The brains were removed and postfixed overnight at 4 °C, followed by a three-day incubation in 30% sucrose in 0.1 M PB. Samples were frozen in Tissue-Tek O.C.T. compound (Sakura Finetek Japan) and coronally sectioned at 60 µm thickness on a cryostat (MICROM HM-560 Thermo Fisher Scientific).

To block non-specific binding, free-floating sections were incubated in a blocking buffer containing 10% Block Ace (Morinaga) in 50 mM tris buffered saline (TBS; pH, 7.4) for 30 minutes. After blocking, they were incubated overnight at 4°C with either goat anti-GFP (1:2000; ab#6673, Abcam) or chicken anti-RFP (1:2000; ab#205402, Abcam) antibodies in 50 mM TBS with 0.2% Triton X. They were then incubated with alexa488 anti-goat (1:1000; Jackson Immune Research) or alexa594 anti-chicken (1:1000; Jackson Immune Research) antibodies in 50 mM TBS with 4′,6-diamidino-2-phenylindole (DAPI).

For viral tracing, additional neuronal nuclei (NeuN) staining was performed using rabbit anti-neuronal nuclei (1:5000; EPR12763, Abcam) with alexa680 anti-rabbit (1:1000; Jackson Immune Research) instead of DAPI staining, using the method described above.

Finally, all sections were mounted on gelatin-coated slides, air-dried, and cover-slipped using Fluoromount-G (Southern Biotechnology Associates).

**Image analysis**

Using a brightfield-fluorescent combined microscope (BZX-2000, KEYENCE), images of the area of interest identified based on the Mouse Brain in Stereotaxic Coordinates (6) were captured and stitched at 20x magnification. To quantitatively analyze the number of immunopositive cells, we used Fiji (ver. 1.53h). The captured images were thresholded using Otsu Thresholding (11), then binarized. The resulting binarized signals were automatically counted as positive signals.

**Statistical analysis**

We used Numpy 1.19.0 for statistical analysis of the fiber photometry and animal movement data derived using Python. All behavioral annotation data obtained by BORIS was statistically analyzed using a student’s t-test or an analysis of variance using GraphPad Prism 9. Differences were considered significant at *p* <0.05*, *p* <0.01**, *p* <0.001***, and *p* <0.0001****. All statistical analyses are summarized in Tables S3 and S4.

**Specific methods for studies 1–4 and the supplemental study**

**Study 1: Fiber photometry recording of MeApd-ERβ^+^ neuronal activity during preference tests**

The neuronal activity of MeApd-ERβ^+^ during preference tests was recorded using fiber photometry methods for 10 minutes in seven naive ERβ-iCre-positive male mice. Each mouse was tested twice, once for the receptivity-based preference of RF vs. XF and once for the sex-based preference of RF vs. IM, on separate days with a fixed order and one-week intervals between tests. We analyzed preference behavior and neuronal activity during the sniffing towards each stimulus mouse using a custom program developed in Python (ver. 3.8.1) and aligned it with the animal behavioral annotation derived by BORIS and DeepLabCut (refer to the above for details of each method). Finally, all animals were perfused, and the fiber insertion sites were confirmed under a microscope (Fig. S7).

**Study 2: Fiber photometry recording of MeApd-ERβ positive neurons towards individually presented social stimuli**

The neuronal activity of MeApd-ERβ^+^ during preference tests was recorded with fiber photometry methods for 10 minutes in five naive ERβ-iCre-positive male mice. Each mouse was tested three times, in the order of RF vs. Empty, XF vs. Empty, and IM vs. Empty, on separate days with one-week intervals between tests. We analyzed preference behaviors and neuronal activity during the sniffing towards each stimulus using a custom program developed in Python (ver. 3.8.1) and aligned them with the animal behavioral annotation derived by BORIS and DeepLabCut (refer to the above for details of each method). Finally, all animals were perfused, and the fiber insertion sites were confirmed under a microscope (Fig. S8).

**Study 3: Effects of DREADD inhibition of MeApd-ERβ^+^ neurons on the RF vs. XF and RF vs. IM preference tests**

A total of 15 naïve ERβ-iCre-positive male mice (eight for hM4Di and seven for the mCherry control) were used in this study. MeApd-ERβ^+^ neurons were inhibited during a 10-minute preference test using DREADD. All mice were tested for preference of RF vs. XF three times, including a baseline control level with saline injection, a CNO injection, and a recovery period with saline injection at a four- to five-day interval.

They were then tested for preference between RF and IM using the same protocol, starting seven days after the last test for RF vs. XF. For automatic quantification of the sniffing behavior of test mice, the Ogawa-Type Social Interaction Test System, which has been previously described (8) was used to track the nose tip and body movements of the animals (refer to the above for details of each method).

**Study 4: Viral tracing of MeApd-ERβ^+^ neurons**

A total of three ERβ-iCre-positive mice were stereotaxically injected with 300 μl of AAV2-Flex-synaptophysin-enhanced GFP into the right MeApd (AP: -1.94 mm, ML: ±2.40 mm, DV: -4.75 mm). After three weeks, the mice were perfused, and viral expression was confirmed by immunohistochemical detection of enhanced GFP protein.

**Study 4: Fiber photometry recording of BNSTp in mice with DREADD inhibition of MeApd-ERβ+ neurons**

A total of six naive ERβ-iCre-positive male mice were used in this study. During the preference tests, the neuronal activity of BNSTp was non-specifically recorded using fiber photometry methods while chemogenetically manipulating the neuronal activity of MeApd-ERβ^+^.

All mice were subjected to two preference tests for RF vs. XF, one week apart, with saline injection in the first test and CNO injection in the second test. One week after the second test, the mice were tested for their preference for RF vs. IM using the same protocol. We analyzed preference behavior and neuronal activity during the sniffing towards each stimulus mouse using a custom program developed in Python (ver. 3.8.1) and aligned it with the animal behavioral annotation derived by BORIS and DeepLabCut (refer to above for details of each method). Finally, all animals were perfused, and the fiber insertion sites were confirmed under a microscope (Fig. S9).

**Supplemental Study: Optogenetic stimulation of** **MeApd-ERβ^+^ neurons during the preference test**

A total of seven naive ERβ-iCre-positive male mice were used in this study. MeApd-ERβ^+^ neurons were optogenetically activated during preference tests. Each mouse underwent two preference tests for RF vs. XF on consecutive days (days 1 and 2). Mice were tested for baseline preference without and with optogenetic stimulation on days 1 and 2, respectively.

Each preference test was conducted for 15 minutes and divided into three 5-minute blocks: pre-stimulation, stimulation, and post-stimulation. On day 2, optogenetic stimulation was manually delivered during the stimulation block while the animal was sniffing the XF through holes on the rounded side of the cylinder. All behavioral annotations were derived using BORIS, and mouse movements were tracked using DeepLabCut (refer to the above for details of each method). Finally, all animals were perfused, and the fiber insertion sites were confirmed under a microscope (Fig. S10).

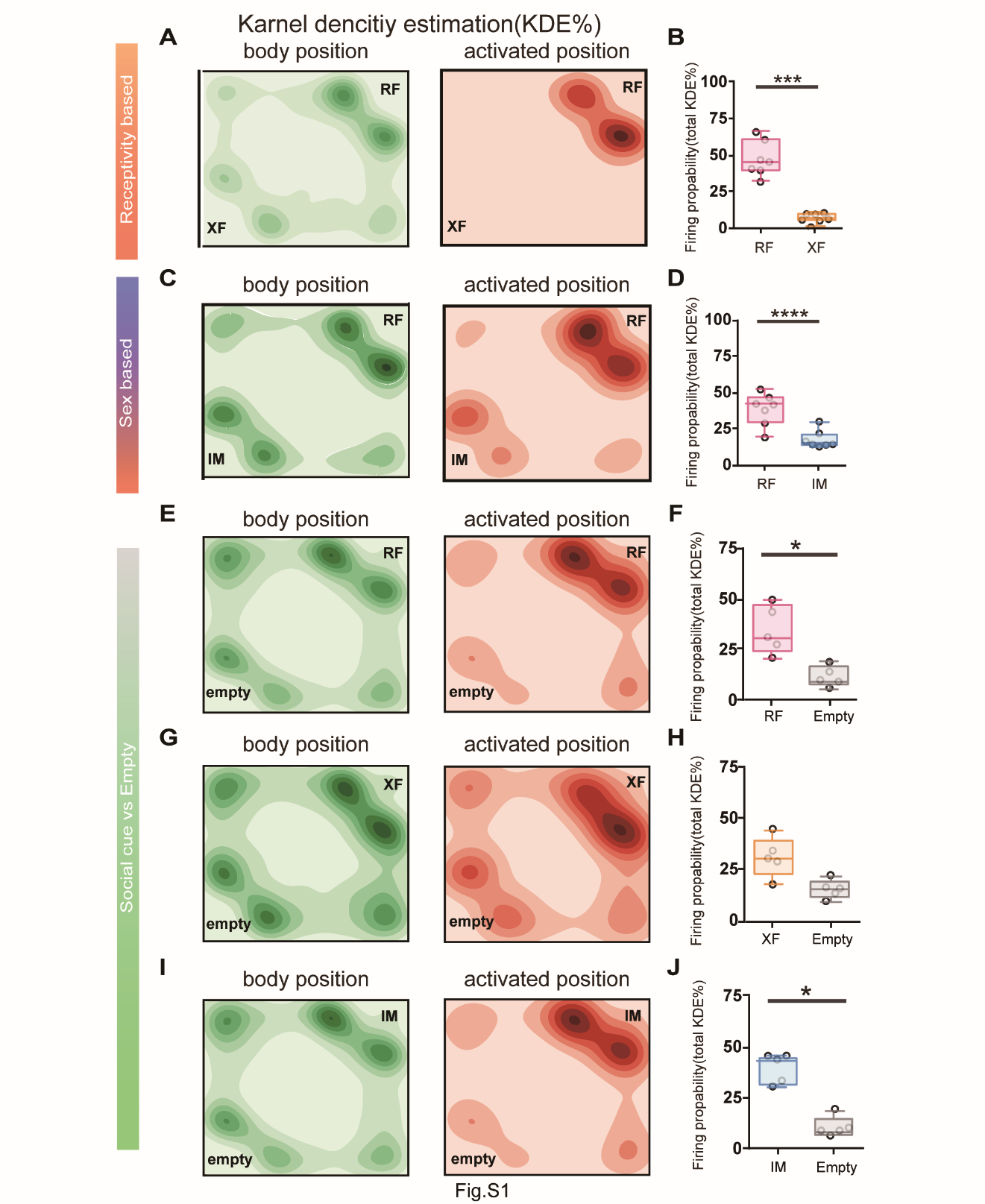
**Supplemental figures**

**Figure S1. Supplemental** **figures for Study 1 and Study 2: Plotting MeApd-ERβ^+^ firing probability by Kernel density estimation during preference test**

(A) Visualization of Kernel density estimation (KDE) in RF vs XF test, shown in a virtual test environment (RF: top right corner, XF bottom left corner). KDE was derived from body positions (Green) and positions with Z > 1 GCaMP7f signals (Red) recorded in 7 animals.

(B) Mean KDE probability score of Z > 1 GCaMP7f signals obtained by summarizing KDE scores in each contact area in RF (Red) vs XF(Orange) preference tests (Mean ± SEM, n = 7, ***: *p* = 0.0008, Student’s t-test).

(C) Visualization of KDE in RF vs IM preference test, shown in a virtual test environment (RF: top right corner, IM bottom left corner). KDE was derived from body positions (Green) and positions with Z > 1 GCaMP7f signals (Red) recorded in 7 animals.

(D) Mean KDE probability score of Z > 1 GCaMP7f signals obtained by summarizing KDE scores in each contact area in RF (Red) vs IM (Blue) preference tests (Mean ± SEM, n = 7, ****: *p* < 0.0001, Student’s t-test).

(E) Visualization of KDE in RF vs Empty test, shown in a virtual test environment (RF: right corner, Empty bottom left corner). KDE was derived from body positions (Green) and positions with Z > 1 GCaMP7f signals (Red) recorded in 5 animals.

(F) Mean KDE probability score of Z > 1 GCaMP7f signals obtained by summarizing KDE scores in each contact area in RF (Red) vs Empty (Gray) preference tests (Mean ± SEM, n = 5, *: *p* = 0.0047 Student’s t-test).

(G) Visualization of KDE in XF vs Empty test shown in a virtual test environment t (XF: right corner, Empty bottom left corner). KDE was derived from body positions (Green) and positions with Z > 1 GCaMP7f signals (Red) recorded in 5 animals.

(H) Mean KDE probability score of Z > 1 GCaMP7f signals obtained by summarizing KDE scores in each contact area in XF (Orange) vs Empty (Gray) tests (Mean ± SEM, n = 5, *p* = 0.1113, Student’s t-test).

(I) Mean KDE probability score of Z > 1 GCaMP7f signals obtained by summarizing KDE scores in each contact area in IM(Blue) vs Empty (Gray) tests (Mean ± SEM, n = 5, *: *p* = 0.0040, Student’s t-test).

(J) Visualization of KDE in IM vs Empty test shown in a virtual test environment (IM: top right corner). KDE was derived from body positions (Green) and positions with Z > 1 GCaMP7f signals (Red) recorded in 5 animals.

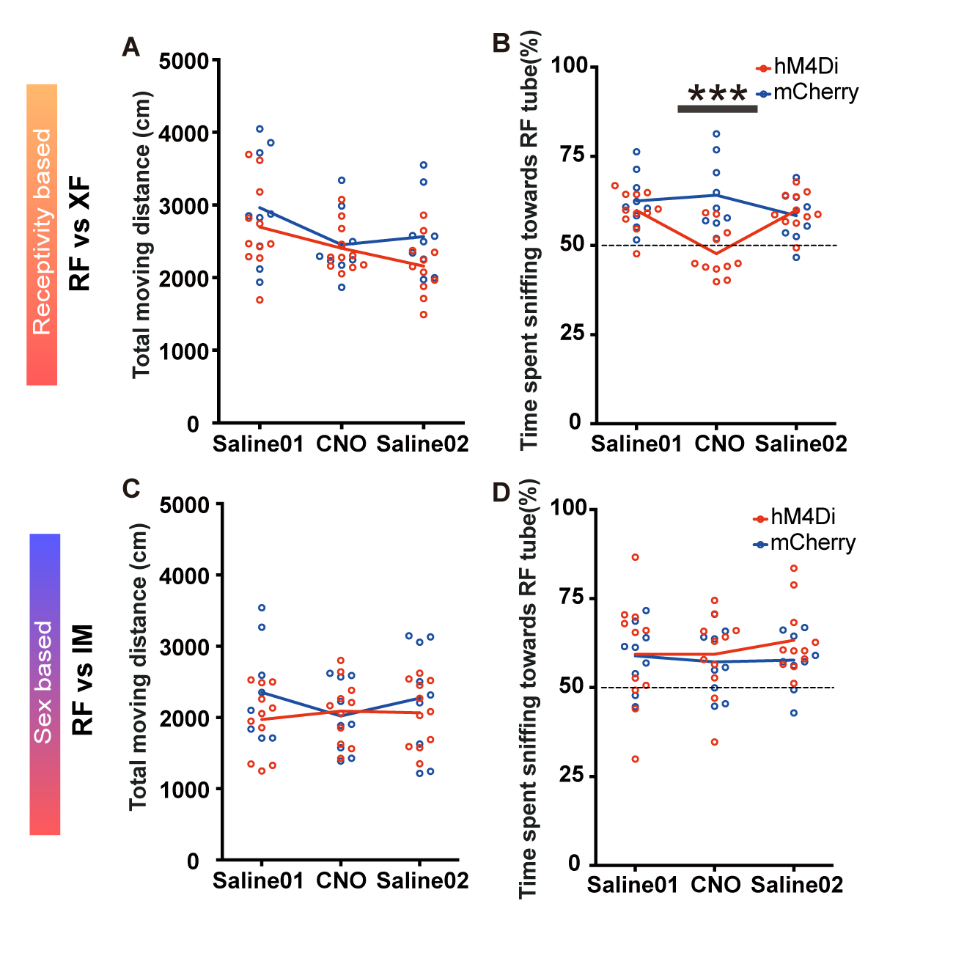

**Figure S2. Supplemental figures for Study 3: DREADD inhibition of MeApd-ERβ^+^ neurons during preference behavior did not affect total animal movement**

(A) Mean total moving distance during RF vs XF tests for hM4Di (Red) and mCherry control (Blue) AAV injected groups (Mean ± SEM, n = 11 for hM4Di and n = 9 for mCherry control, ns, Bonferroni’s test).

(B) Mean percent (%) of time spent sniffing towards RF cylinder during RF vs XF tests for hM4Di (Red) and mCherry control (Blue) groups. Values obtained by dividing RF sniffing duration by total sniffing duration to RF and XF. (Mean ± SEM, n = 11 for hM4Di and n = 9 for mCherry control, ****: *p* < 0.0001, Bonferroni’s test).

(C) Mean total moving distance during RF vs IM tests for hM4Di (Red) and mCherry control (Blue) AAV injected groups (Mean ± SEM, n = 11 for hM4Di and n = 9 for mCherry control, ns, Bonferroni’s test).

(D) Mean percent (%) of time spent sniffing towards RF cylinder during RF vs IM tests for hM4Di (Red) and mCherry control (Blue) groups. Values obtained by dividing RF sniffing duration by total sniffing duration to RF and IM. (Mean ± SEM, n = 11 for hM4Di and n = 9 for mCherry control, ns, Bonferroni’s test).

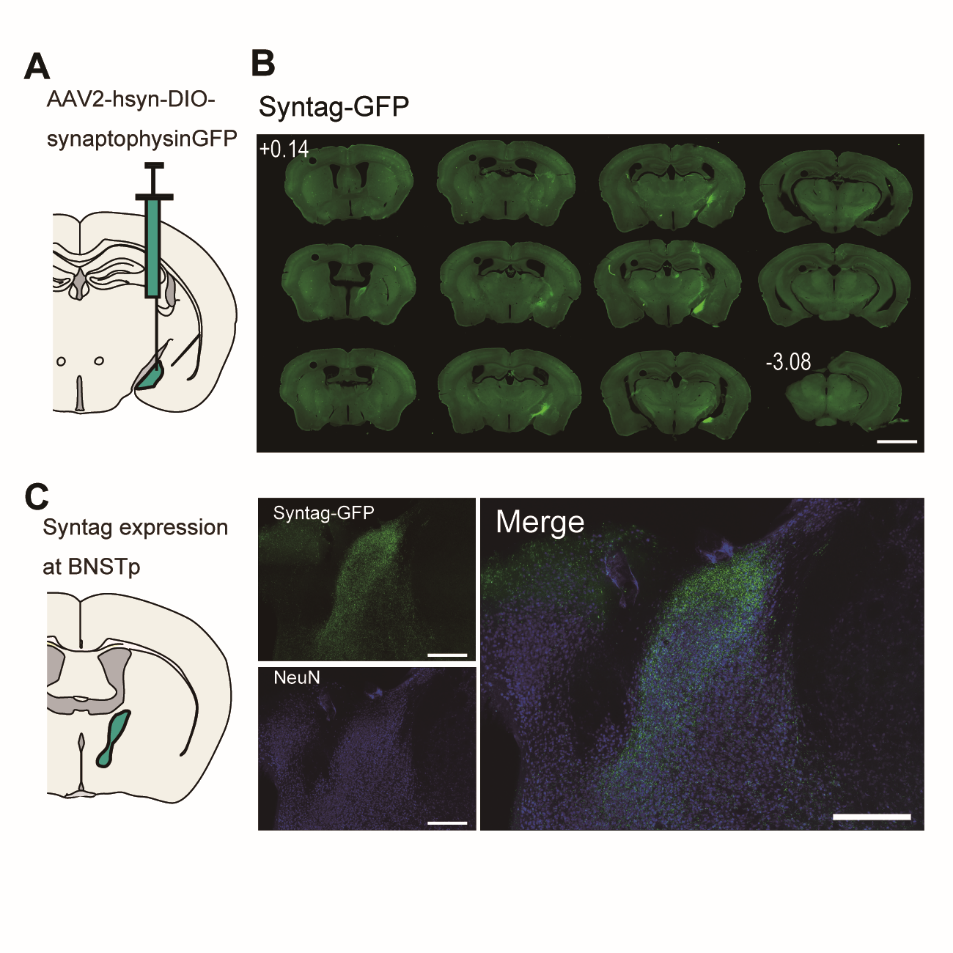

**Figure S3. Supplemental figures for Study 4:** **Viral tracing of MeApd-ERβ^+^ neurons**

(A) Schematic diagram of unilateral AAV injection for tracing MeApd-ERβ^+^ neurons.

(B) Representative low magnification image showing terminals (GFP) of MeApd-ERβ^+^ neurons across Bregma 0.14mm to -3.08mm (slice 0.24mm each, scale bar = 2.5mm).

(C) Representative image showing expression of terminals (GFP, left top) of MeApd-ERβ^+^ neurons in neurons stained with NeuN (Blue, left bottom) in BNSTp as shown in merged image (right) (scale bar = 500um).

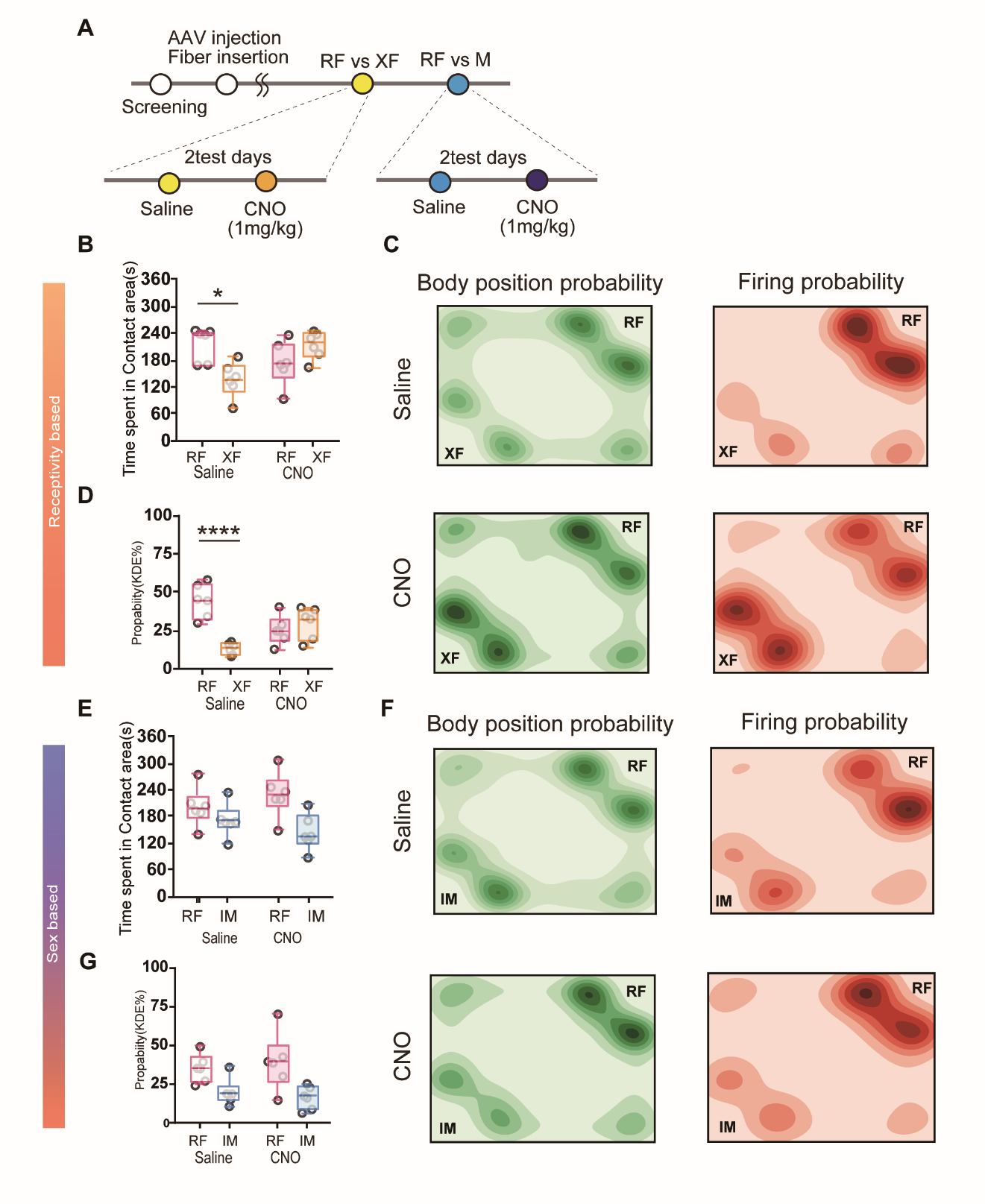

**Figure S4. Supplemental figures for Study 4:** **Effects of DREADD inhibition of MeApd-ERβ^+^ neurons on preference behavior and BNSTp firing probability**

(A)Timeline for preference tests with DREADD inhibition. Male mice were injected with AAV and tested for receptivity-based and sex-based preference in two consecutive days each with either saline (day 1) or CNO (day 2; 1 mg/kg) injections(right).

(B) Duration of time (sec) spent in contact area during RF (Red) vs XF(Orange) tests in saline and CNO.

(C) Visualization of KDE shown in virtual test environment (RF: top right corner XF: bottom left corner). KDE was derived from body positions (Green) and positions with Z > 1 GCaMP7f signals (Red) recorded in 6 animals. Upper row represents KDE during saline administration, and bottom row represents KDE during CNO administration.

(D) Mean KDE probability of Z > 1 GCaMP7f signals obtained by summarizing KDE scores in each contact area in RF (Red) vs XF(Orange) tests (Mean ± SEM, n = 6, ****: *p* < 0.0001, Bonferroni’s test).

(E) Duration of time spent in contact area during RF (Red) vs IM(Blue) tests in each saline and CNO injected session (Mean ± SEM, n=6, ns., Bonferroni’s test).

(G) Visualization of KDE shown in a virtual test environment (RF: top right corner IM: bottom left corner). KDE was derived from body positions (Green) and positions with Z > 1 GCaMP7f signals (Red) recorded in 6 animals. Upper row represents KDE during saline administration, and bottom row represents KDE during CNO administration.

(F) Mean KDE probability of Z > 1 GCaMP7f signals obtained by summarizing KDE scores in each contact area in RF (Red) vs IM (Blue) tests (Mean ± SEM, n = 6, ns, Bonferroni’s test).

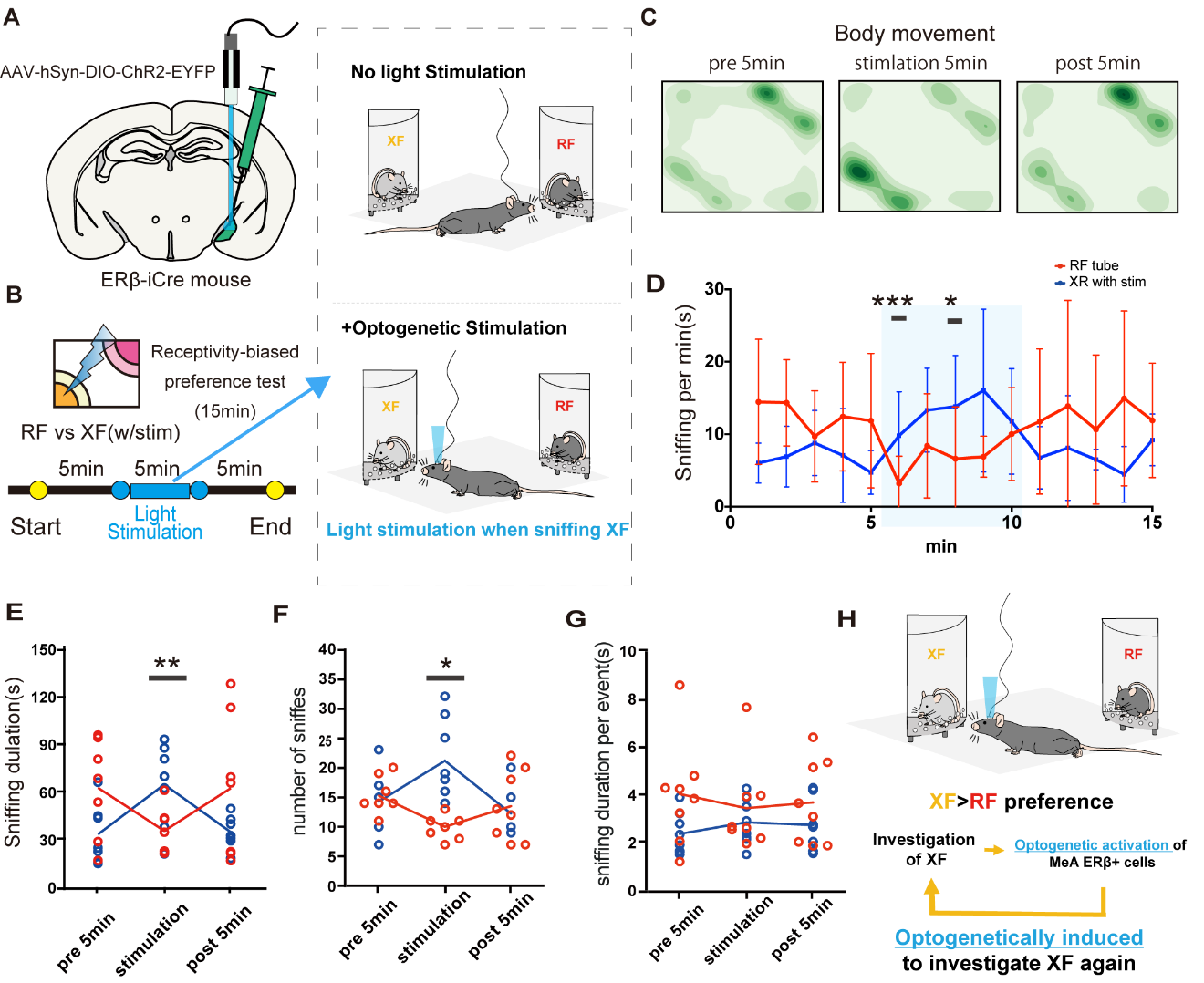

Figure S5. Figures for Supplemental Study: Effects of optogenetic stimulation of MeApd-ERβ^+^ neurons on the levels of sniffing behavior

(A) Schematic diagram of AAV injection and fiber implantation in the MeApd and receptive female (RF) vs non-receptive female (XF) preference test with fiber-photometry recordings.

(B) Schematic diagram and timeline of fiber-photometry recordings with optogenetic stimulation. Male mice were introduced in a receptivity-based (RF vs XF) preference test environment for 15 min. Each test was divided into three 5min blocks (pre-stimulation, stimulation, post-stimulation). Optogenetic stimulation was delivered manually during the stimulation block while the animal was sniffing XF.

(C) Mean heat map presentation of body positions of male mice (n = 5) during the first 5 min (pre-stimulation), middle (stimulation) and last 5 min (post-stimulation).

(D) Mean sniffing duration per minute towards RF (Red) and XF (Blue) during 15 min tests (Mean ± SEM, n = 5, *, *p* < 0.05, ***: *p* < 0.001, Bonferroni’s test).

(E) Mean sniffing duration in each 5 min block during RF(Red) vs XF(Blue) tests (Mean ± SEM, n = 5, *: *p* < 0.05, **: *p* < 0.01, Bonferroni’s test).

(F) Mean number of sniffing events in each 5 min block during RF(Red) vs XF(Blue) tests (Mean ± SEM, n = 5, *: *p* < 0.05, Bonferroni’s test).

(G) Mean sniffing duration per sniffing event in each 5 min block during RF(Red) vs XF(Blue) tests (Mean ± SEM, n = 5, ns, Bonferroni’s test).

(H) Schematic diagram demonstrating possible mechanisms of MeApd-ERβ^+^ neuronal excitation inducible sniffing and preference to RF.

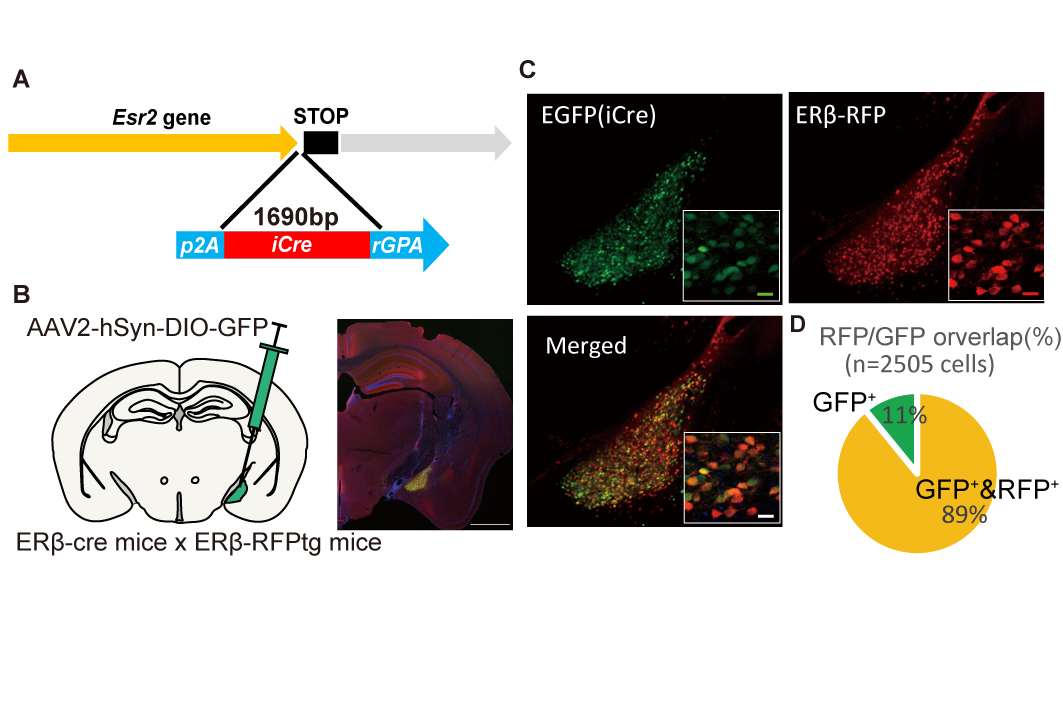

**Figure S6. Generation and verification of ERβ-iCre mice**

(A)Schematic diagram for generation of ERβ-iCre mice using CRISPR-Cas9. The 2A-iCre-rGPA sequence was knocked in right before the Stop codon of *Esr2* gene.

(B)Schematic diagram of AAV injection for Cre-dependent induction of GFP to the MeA in ERβ-iCre and ERβ-RFPtg double positive mice (left). EGFP and RFP expressing cells were localized in the MeApd (right), (Scale bar = 1 mm).

(C)Representative high magnification images showing iCre-dependently induced EGFP positive cells (top left), ERβ-RFP positive cells (top right), and merged image (bottom) in the MeApd (Scale bar = 10 μm).

(D)Percentage of RFP positive cells co-expressing EGFP out of a total of 2505 EGFP positive cells (accumulated data from 3 mice) in MeApd.

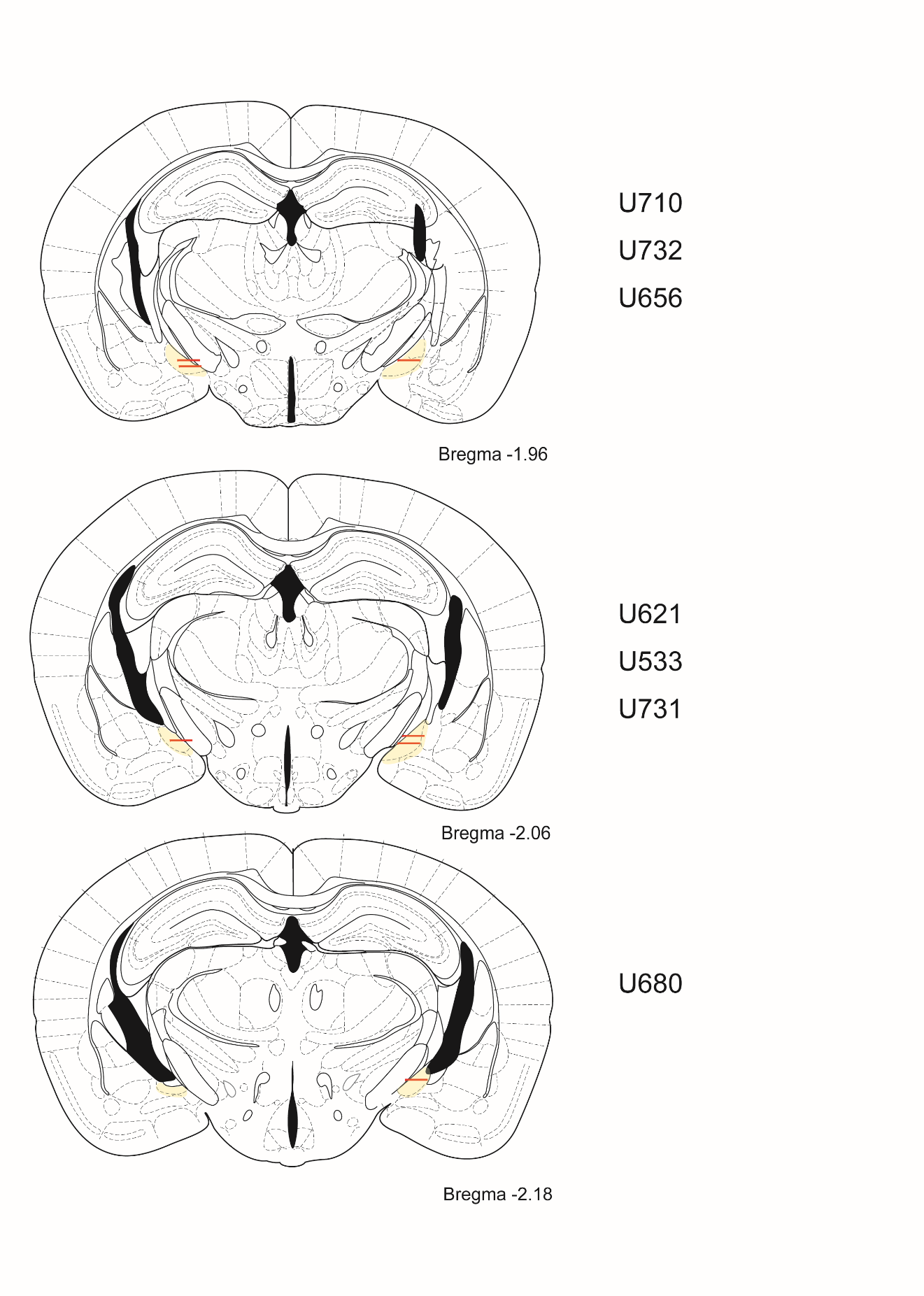
Figure S7. Fiber insertion sites for animals used in Study 1.

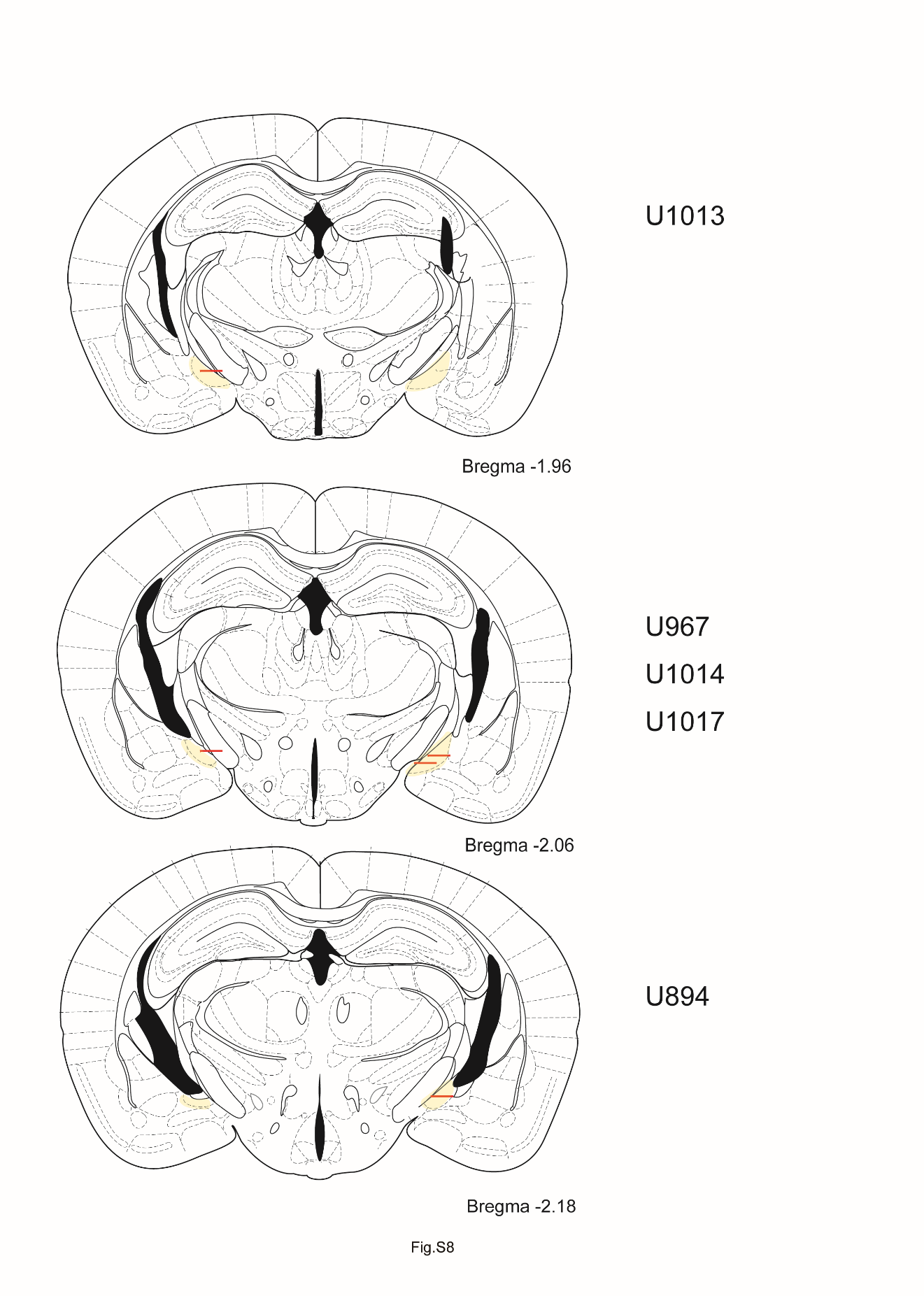
Figure S8. Fiber insertion sites for animals used in Study 2.

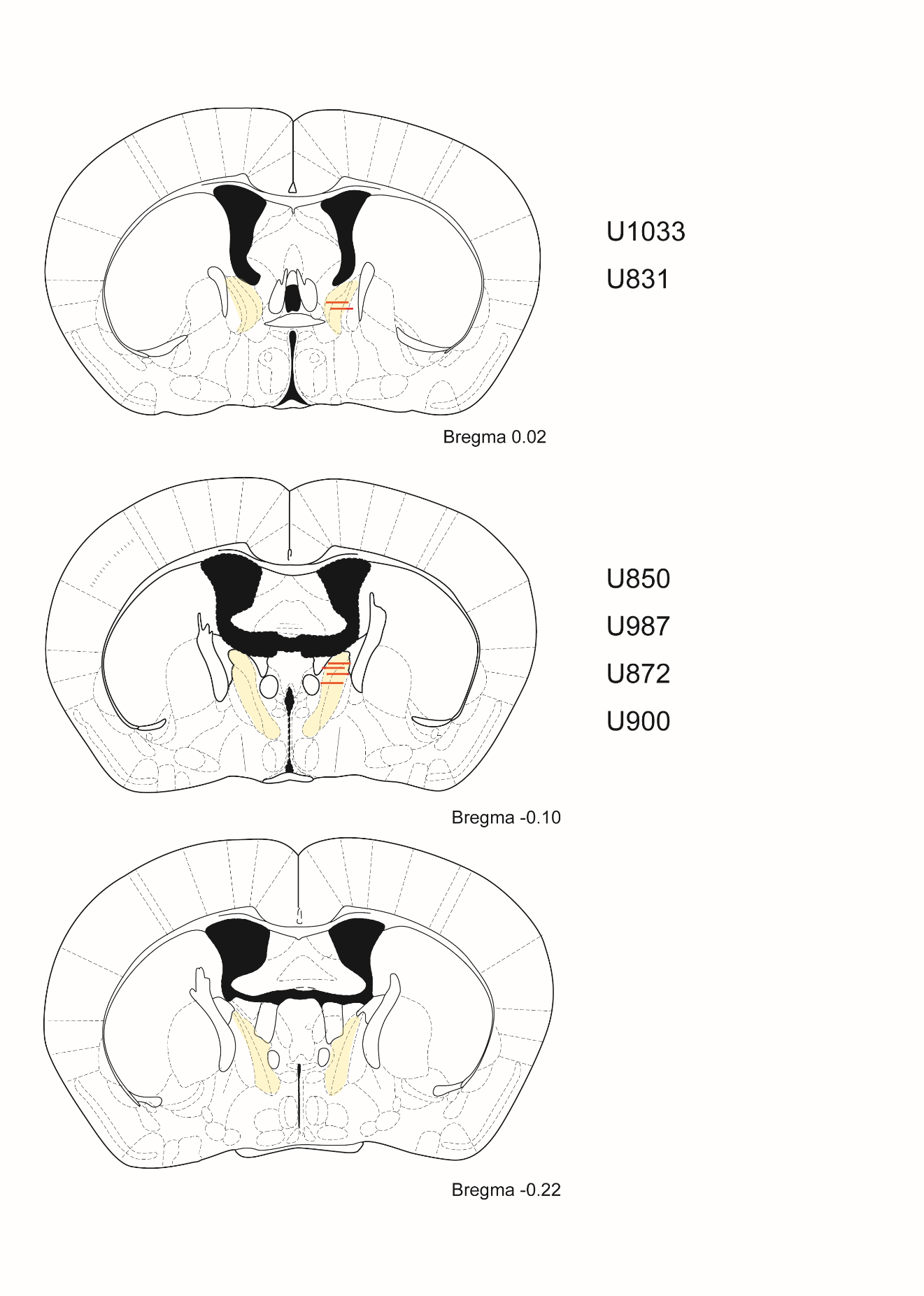
Figure S9. Fiber insertion sites for animals used in Study 4.

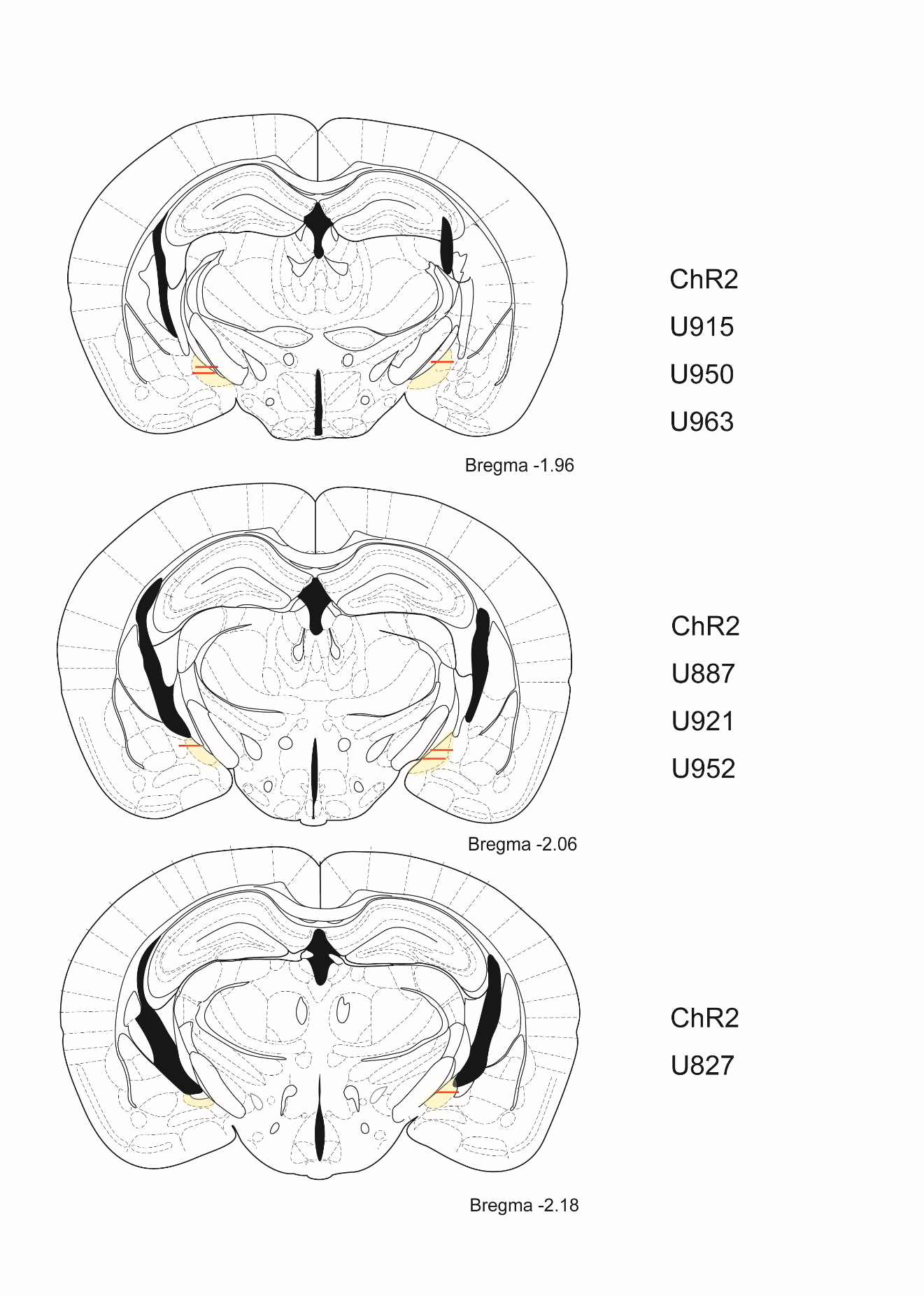
Figure S10. Fiber insertion sites for animals used in Supplemental Study.

Supplemental tables

Table S1. Primers used for screening ERβ-iCre mice

| sequence | description |
| --- | --- |
| GTCCCTGGTGATGAGGAGAA | iCre detect F |
| ATCAGCATTCTCCCACCATC | iCre detect R |
| TACAGCTTGGTGATGAGGTTTTGCTCTT | EsrCre detect 5F |
| AGATCCATCTCTCCACCAGCTTGGTAAC | EsrCre detect 5R |
| ACAGACAGGAGCATCTTCCA | iCre seq R |
| GAGGATGTGAGGGACTACCTCCTGTACC | EsrCre detect 3F |
| ACACATTTTGTGAATTTGCCATGTTCCT | EsrCre detect 3R |
| AGTTCATCAAGCCCATCCTG | Cas9 detection F |
| GAAGTTTCTGTTGGCGAAGC | Cas9 detection R |
| TTGCCGGGAAGCTAGAGTAA | Amp detection F |
| TTTGCCTTCCTGTTTTTGCT | Amp detection R |

Table S2. AAV related plasmids used in this study

| Plasmid name | Source | Reference |
| --- | --- | --- |
| pGp-AAV-syn-jGCaMP7f-WPRE | pGP-AAV-syn-jGCaMP7f-WPRE was a gift from Douglas Kim & GENIE Project (Addgene plasmid # 104488 ; <http://n>2t.net/addgene:104488 ; RRID:Addgene_104488) | (12) |
| pAAV-hSyn-DIO-EGFP | pAAV-hSyn-DIO-EGFP was a gift from Bryan Roth (Addgene plasmid # 50457 ; <http://n>2t.net/addgene:50457 ; RRID:Addgene_50457) | unpublished |
| pAAV-hSyn-DIO-mCherry | pAAV-hSyn-DIO-mCherry was a gift from Bryan Roth (Addgene plasmid # 50459 ; <http://n>2t.net/addgene:50459 ; RRID:Addgene_50459) | (13) |
| pAAV-hSyn-DIO-hM4D(Gi)-mCherry | pAAV-hSyn-DIO-hM4D(Gi)-mCherry was a gift from Bryan Roth (Addgene plasmid # 44362 ; <http://n>2t.net/addgene:44362 ; RRID:Addgene_44362) | (13) |
| pAAV-Ef1a-DIO hChR2(E123A)-EYFP | pAAV-Ef1a-DIO hChR2(E123A)-EYFP was a gift from Karl Deisseroth (Addgene plasmid # 35507 ; <http://n>2t.net/addgene:35507 ; RRID:Addgene_35507) | (14) |
| pAAV-FLEX-Synaptophysin-GFP | pAAV FLEX Synaptophysin GFP was a gift from Matthew Nolan (Addgene plasmid # 137188 ; <http://n>2t.net/addgene:137188 ; RRID:Addgene_137188) | (15) |

Table S3. Statistical summary for the main figures

| Experiment | Figure(s) | Statistical details |
| --- | --- | --- |
| Fiber photometry GcaMP7f imaging of MeA ERβ positive cells during RF vs XF preference test | Figure 1E  Time spent in the contact area | Student’s t-test n=7, *t*=5.428 *df*=6 *p*=0.0008*** |
|  | Figure 1F  Sniffing duration | Student’s t-test n=7, *t*=6.31 *df*=6 *p*=0.0004*** |
|  | Figure 1G  Mean zFn | Student’s t-test n=7, *t*=8.293 *df*=6 *p*<0.0001**** |
| Fiber photometry GcaMP7f imaging of MeA ERβ positive cells during RF vs IM preference test | Figure 1L  Time spent in the contact area | Student’s t-test n=7, *t*=1.874 *df*=6 *p*=0.0550 |
|  | Figure 1M  Sniffing duration | Student’s t-test n=7, *t*=3.591 *df*=6 *p*=0.0057*** |
|  | Figure 1O  Mean zFn | Student’s t-test n=7, *t*=3.114 *df*=6 *p*=0.0104* |

| Experiment | Figure(s) | Statistical details |
| --- | --- | --- |
| Fiber photometry GcaMP7f imaging of MeA ERβ positive cells  during RF vs empty preference test | Figure 2A  Time spent in the contact area | Student’s t-test n=5, *t*=2.009 *df*=4 *p*=0.0576 |
|  | Figure 2C  Mean peak zFn | Student’s t-test n=5, *t*=3.511 *df*=4 *p*=0.0123* |
| Fiber photometry GcaMP7f imaging of MeA ERβ positive cells  during XF vs empty preference test | Figure 2G  Time spent in the contact area | Student’s t-test n=5, *t*=3.480 *df*=4 *p*=0.0127* |
|  | Figure 2H  Mean peak zFn | Student’s t-test n=5, *t*=1.576 *df*=4 *p*=0.0951 |
| Fiber photometry GcaMP7f imaging of MeA ERβ positive cells  during IM vs empty preference test | Figure 2I  Time spent in the contact area | Student’s t-test n=5, *t*=8.642 *df*=4 *p*=0.0005*** |
|  | Figure 2K  Mean peak zFn | Student’s t-test n=5, *t*=3.516 *df*=4 *p*=0.0123* |

| Experiment | Figure(s) | Statistical details |
| --- | --- | --- |
| DREADD inhibition of MeA ERβ positive cells during RF vs XF preference test  (Stimulation at the XF side) | Figure 3D  Time spent in the RF side (%) | Repeated measures ANOVA,  Drug vs Virus  Interaction (drug x virus) *F* _(2,36)_ =7.196 *p*=0.0024***  post-hoc Bonferroni’s multiple comparisons test  mCherry(n=9) – hM4Di(n=11) x drug  CNO: Adjusted *p* value<0.0001**** |
| DREADD inhibition of MeA ERβ positive cells during RF vs IM preference test  (Stimulation at the IM side) | Figure 3E  Time spent in the RF side (%) | Repeated measures ANOVA,  Drug vs Virus  Interaction (drug x virus) *F* _(2,36)_ =0.2953 *p=*0.7535 *ns* |

| Experiment | Figure(s) | Statistical details |
| --- | --- | --- |
| Fiber photometry GcaMP7f imaging of BNSTp cells and CNO manipulation of MeA ERβ positive cells  during RF vs XF preference test | Figure 4F  Mean peak zFn | Repeated measures ANOVA,  drug vs cue(n=6)  Interaction (drug x cue) *F* _(1,20)_ =13.95 *p<*0.0001****  post-hoc Bonferroni’s multiple comparisons test  RF – XF x drug  Saline: Adjusted *p<*0.0007***  CNO: Adjusted *p* value>0.9999  Saline – CNO x cue  RF: Adjusted *p* value=0.0110**  XF: Adjusted *p* value=0.0035** |
| , Fiber photometry GcaMP7f imaging of BNSTp cells and CNO manipulation of MeA ERβ positive cells  during RF vs IM preference test | Figure 4I  Mean peak zFn | Repeated measures ANOVA,  drug vs cue(n=6)  Interaction (drug x cue) *F* _(1,20)_ =0.3787 *p=*0.5452  Cue *F* _(1,20)_ =9.162 *p=*0.0067** |

Table S4. Statistical summary for supplemental figures

| Experiment | Figure(s) | Statistical details |
| --- | --- | --- |
| Fiber photometry GcaMP7f imaging of MeA ERβ positive cells  during RF vs XF preference test | Figure S1B  Firing probability driven by KDE analysis | Student’s t-test n=7, *t*=5,462 *df*=6 *p*=0.0008*** |
| Fiber photometry GcaMP7f imaging of MeA ERβ positive cells  during RF vs IM preference test | Figure S1D  Firing probability driven by KDE analysis | Student’s t-test n=7, *t*=7.545 *df*=6 *p*<0.0001*** |
| Fiber photometry GcaMP7f imaging of MeA ERβ positive cells  during RF vs empty preference test | Figure S1F  Firing probability driven by KDE analysis | Student’s t-test n=5, *t*=4.693 *df*=4 *p*=0.0047* |
| Fiber photometry GcaMP7f imaging of MeA ERβ positive cells  during XF vs empty preference test | Figure S1H  Firing probability driven by KDE analysis | Student’s t-test n=5, *t*=1.442 *df*=4 *p*=0.1113 |
| Fiber photometry GcaMP7f imaging of MeA ERβ positive cells  during IM vs empty preference test | Figure S1J  Firing probability driven by KDE analysis | Student’s t-test n=5, *t*=4.897 *df*=4 *p*=0.0040** |

| Experiment | Figure(s) | Statistical details |
| --- | --- | --- |
| DREADD inhibition of MeA ERβ positive cells during RF vs XF preference test  (Stimulation at the XF side) | FigureS2A  Total moving distance | Repeated measures ANOVA,  Drug vs Virus  Interaction (drug x virus) *F* _(2,36)_ =0.7302 *p*=0.4999 *ns* |
|  | Figure S2B  Time spent sniffing towards RF tube (%) | Repeated measures ANOVA,  Drug vs Virus  Interaction (drug x virus) *F* _(2,36)_ =7.244 *p*=0.0023***  post-hoc Bonferroni’s multiple comparisons test  mCherry(n=9) - hM4Di(n=11) x drug  CNO: Adjusted *p* value<0.0001**** |
| DREADD inhibition of MeA ERβ positive cells during RF vs IM preference test  (Stimulation at the IM side) | Figure S2C  Total moving distance | Repeated measures ANOVA,  Drug vs Virus  Interaction (drug x virus) *F* _(2,36)_ =1.899 *p=*0.1645 *ns* |
|  | Figure S2D  Time spent sniffing towards RF tube (%) | Repeated measures ANOVA,  Drug vs Virus  Interaction (drug x virus) *F* _(2,36)_ =1.612 *p=*0.2136 *ns* |

| Experiment | Figure(s) | Statistical details |
| --- | --- | --- |
| Optogenetic stimulation of MeA ERβ positive cells during RF vs XF preference test  (Stimulation at the XF side) | Figure S4B  Time spent in contact aria (every min) | Repeated measures ANOVA,  Time vs social cue (RF vs XF)  Interaction (time x cue) *F* _(14,112)_ =5.976 *p*<0.0001****  post-hoc Bonferroni’s multiple comparisons test(n=5)  RF – XF (min from test)  6min: Adjusted *p* value=0.0004***  7min: Adjusted *p* value=0.2302  8min: Adjusted *p* value=0.0449*  9min: Adjusted *p* value=0.0672 |
|  | Figure S4D  Time spent sniffing (min) | Repeated measures ANOVA,  Time vs social cue (RF vs XF)  Interaction (time x cue) *F* _(2,16)_ =18.30 *p*<0.0001****  post-hoc Bonferroni’s multiple comparisons test(n=5)  Sniff RF-XF  pre stim: Adjusted *p* value=0.0281*  stim: Adjusted *p* value=0.0064**  post stim: Adjusted *p* value=0.261 |
|  | Figure S4E  Number of sniffing | Repeated measures ANOVA,  Time vs social cue (RF vs XF)  Interaction (time x cue) *F* _(2,16)_ =20.89 *p*<0.0001****  post-hoc Bonferroni’s multiple comparisons test(n=5)  Sniff RF-XF  pre stim: Adjusted *p* value=0.9926  stim: Adjusted *p* value=0.0142*  post stim: Adjusted *p* value=0.8573 |
|  | Figure S4G  Duration of individual sniffing event | Repeated measures ANOVA,  Time vs social cue (RF vs XF)  Interaction (time x cue) *F* _(2,16)_ =1.966 *p=*0.1723 |

| Experiment | Figure(s) | Statistical details |
| --- | --- | --- |
| Fiber photometry GCaMP7f imaging of BNSTp cells and CNO manipulation of MeA ERβ positive cells  during RF vs XF preference test | Figure S5C  Time spent in the contact area | Repeated measures ANOVA,  drug vs cue(n=6)  Interaction (drug x cue) *F* _(1,20)_ =13.64 *p*=0.0014**  post-hoc Bonferroni’s multiple comparisons test  RF – XF x drug  Saline: Adjusted *p* value=0.0127*  CNO: Adjusted *p* value=0.6331  Saline – CNO x cue  RF: Adjusted *p* value=0.4480  XF: Adjusted *p* value=0.0195* |
|  | Figure S5D  Firing probability driven by KDE analysis | Repeated measures ANOVA,  drug vs cue(n=6)  Interaction (drug x cue) *F* _(1,20)_ =23.80 *p<*0.0001****  post-hoc Bonferroni’s multiple comparisons test  RF – XF x drug  Saline: Adjusted *p<*0.0001****  CNO: Adjusted *p* value>0.9999  Saline – CNO x cue  RF: Adjusted *p* value=0.009*  XF: Adjusted *p* value=0.0257* |
| Fiber photometry GCaMP7f imaging of BNSTp cells and CNO manipulation of MeA ERβ positive cells  during RF vs IM preference test | Figure S5G  Time spent in the contact area | Repeated measures ANOVA,  drug vs cue(n=6)  Interaction (drug x cue) *F* _(1,20)_ =2.375 *p=*0.1389  Cue *F* _(1,20)_ =10.90 *p=*0.0036** |
|  | Figure S5H  Firing probability driven by KDE analysis | Repeated measures ANOVA,  drug vs cue(n=6)  Interaction (drug x cue) *F* _(1,20)_ =0.6713 *p=*0.4223  Cue *F* _(1,20)_ =16.53 *p=*0.0006** |

SI References

1.	G. R. Terrell, D. W. Scott, Variable kernel density estimation. *Ann. Stat.* **20**, 1236–1265 (1992).

2.	Y. Hasegawa, *et al.*, Generation of CRISPR/Cas9-mediated bicistronic knock-in *ins1-cre* driver mice. *Exp. Anim.* **65**, 319–327 (2016).

3.	J. G. Lemmen, *et al.*, Expression of estrogen receptor alpha and beta during mouse embryogenesis. *Mech. Dev.* **81**, 163–167 (1999).

4.	X. Fan, M. Warner, J.-Å. Gustafsson, Estrogen receptor β expression in the embryonic brain regulates development of calretinin-immunoreactive GABAergic interneurons. *Proc. Natl. Acad. Sci.* **103**, 19338–19343 (2006).

5.	S. Sagoshi, *et al.*, Detection and characterization of estrogen receptor beta expression in the brain with newly developed transgenic mice. *Neuroscience* **438**, 182–197 (2020).

6. ,	Paxinos, G. & Franklin, K. B. J. (2001) The mouse brain in stereotaxic coordinates (Academic, San Diego).

7.	M. C. Tsuda, S. Ogawa, Long-lasting consequences of neonatal maternal separation on social behaviors in ovariectomized female mice. *PLOS ONE* **7**, e33028 (2012).

8.	M. Nakata, *et al.*, Effects of prepubertal or adult site-specific knockdown of estrogen receptor β in the medial preoptic area and medial amygdala on social behaviors in male mice. *eNeuro* **3**, ENEURO.0155-15.2016 (2016).

9.	O. Friard, M. Gamba, BORIS: a free, versatile open-source event-logging software for video/audio coding and live observations. *Methods Ecol. Evol.* **7**, 1325–1330 (2016).

10.	A. Mathis, *et al.*, DeepLabCut: markerless pose estimation of user-defined body parts with deep learning. *Nat. Neurosci.* **21**, 1281–1289 (2018).

11.	N. Otsu, A threshold selection method from gray-level histograms. *IEEE Trans. Syst. Man Cybern.* **9**, 62–66 (1979).

12.	H. Dana, *et al.*, High-performance calcium sensors for imaging activity in neuronal populations and microcompartments. *Nat. Methods* **16**, 649–657 (2019).

13.	M. J. Krashes, *et al.*, Rapid, reversible activation of AgRP neurons drives feeding behavior in mice. *J. Clin. Invest.* **121**, 1424–1428 (2011).

14.	J. Mattis, *et al.*, Principles for applying optogenetic tools derived from direct comparative analysis of microbial opsins. *Nat. Methods* **9**, 159–172 (2011).

15.	G. Sürmeli, *et al.*, Molecularly defined circuitry reveals input-output segregation in deep layers of the medial entorhinal cortex. *Neuron* **92**, 929 (2016).
